# JNK-dependent intestinal barrier failure disrupts host-microbe homeostasis during tumorigenesis

**DOI:** 10.1101/719468

**Authors:** Jun Zhou, Michael Boutros

## Abstract

In all animals, the intestinal epithelium forms a tight barrier to the environment. The epithelium regulates the absorption of nutrients, mounts immune responses and prevents systemic infections. Here, we investigate the consequences of tumorigenesis on the microbiome using a *Drosophila* intestinal tumor model. We show that upon loss-of BMP signaling, tumors lead to aberrant activation of JNK signaling, followed by intestinal barrier dysfunction and commensal imbalance. In turn, the dysbiotic microbiome triggers a regenerative response and stimulates tumor growth. We find that inhibiting JNK signaling or depletion of the microbiome restores barrier function of the intestinal epithelium, leading to a reestablishment of host-microbe homeostasis, and organismic lifespan extension. Our experiments identify a JNK-dependent feedback amplification loop between intestinal tumors and the microbiome. They also highlight the importance of controlling the activity level of JNK signaling to maintain epithelial barrier function and host-microbe homeostasis.

## INTRODUCTION

The intestinal epithelium forms a physical barrier that allows selective absorption of nutrients, prevents invasion of pathogens and mounts immune responses. In *Drosophila*, the intestinal epithelium contains highly proliferative intestinal stem cells (ISCs) that produce progenitor cells called enteroblast (EBs). EBs differentiate into either enterocytes (ECs) or enteroendocrine cells (EEs) to ensure homeostatic turnover of the tissue (reviewed in Miguel-Aliaga *et al*, 2018). Several well conserved signaling pathways, including Notch, BMP, Wnt, JAK/STAT and Ras, are required for maintaining intestinal homeostasis and their dysregulation can lead to tumor formation in the *Drosophila* intestine (Guo *et al*, 2013; Patel *et al*, 2015; Ragab *et al*, 2011; Suijkerbuijk *et al*, 2016; Zhai *et al*, 2015; Cordero *et al*, 2012; Parvy *et al*, 2018; Siudeja *et al*, 2015).

Across the animal kingdom, the intestinal epithelium is exposed to a dynamic and metabolically complex bacterial community. Host-microbiome interactions play important roles in maintaining organismal homeostasis, influence the development of the host immune system and help to digest and absorb nutrients (Buchon *et al*, 2013a; Round & Mazmanian, 2009; Hooper *et al*, 2012). While it has become evident that the microbiome is important for organismal health, how changes in host-microbiome interactions contribute to tumorigenesis remains controversially discussed (Tilg *et al*, 2018).

Due to the low diversity of its intestinal microbiota and powerful genetic tools, *Drosophila* has become an important model system to study host-microbe interactions. In *Drosophila*, the microbiota has been reported to be involved in host development (Storelli *et al*, 2011; Yamada *et al*, 2015; Sannino *et al*, 2018; Martino *et al*, 2018; Reedy *et al*, 2019; Kamareddine *et al*, 2018), influencing lifespan (Guo *et al*, 2014; Iatsenko *et al*, 2018; Li *et al*, 2016; Obata *et al*, 2018) and affecting animal behavior and disease (Erkosar *et al*, 2015; Jones *et al*, 2015; Sansone *et al*, 2015; Charroux *et al*, 2018; Fischer *et al*, 2017).

In this study, we created an inducible tumor model by tissue-specific depletion of BMP signaling pathway components and investigated its effect on host-microbe interactions. We show that intestinal tumors lead to epithelial barrier dysfunction, which in turn promotes commensal dysbiosis with an increase in microbial load and a decrease in microbial diversity. Dysbiosis further stimulates intestinal regeneration and tumor growth. Mechanistically, we provide evidence that tumor related JNK hyperactivation leads to barrier loss, dysbiosis and increased mortality. In contrast, either JNK inhibition or microbiota depletion reverses these phenotypes. Thus, our findings support a model whereby the gut microbiome is neither driver or passenger, but acts as part of a self-enforcing feedback loop that promotes further intestinal barrier dysfunction and tumorigenesis through the activation of JNK.

## RESULTS

### The intestinal microbiome promotes BMP-induced tumorigenesis

Dpp/BMP signaling in *Drosophila* is activated by dimers of Dpp ligands that bind to type I (Thickveins; Tkv) and type II (Punt) receptors. Activated Tkv, in turn, phosphorylates the *Drosophila* Smad Mother-against-Dpp (Mad), which interacts with the co-Smad Medea (Med) and Schnurri (Shn). Inactivation of BMP pathway components in the *Drosophila* midgut leads to intestinal tumor phenotypes similar to juvenile polyposis syndrome (JPS) (Guo *et al*, 2013). Previous studies reported multiple roles of Dpp/BMP signaling in intestinal homeostasis in *Drosophila*, including ISC self-renewal, EB to EC differentiation, copper cell differentiation and EC survival (Guo *et al*, 2013; Li *et al*, 2013, 2016; Tian & Jiang, 2014; Tian *et al*, 2017; Zhou *et al*, 2015).

Using the inducible escargot-Flip Out (esgts F/O) system to generate GFP-marked mosaic clones (Jiang *et al*, 2009), we induced the expression of hairpin RNAi constructs targeting BMP-signaling components in ISCs, EBs (escargot+ [esg+] cells) and all their progeny. In these clones, RNAi-expressing cells and their progeny are marked with GFP, while already existing differentiated cells do not express the RNAi construct and remain GFP negative (Fig. 1A). Upon Med or Shn RNAi, we found disorganized, multilayered epithelial clones as compared to a single epithelial layer in controls (Fig 1B-D). The effect of the RNAi was confirmed with wing-specific driver using Nubbin-Gal4, which showed expected BMP loss of function related phenotypes (Fig. S1A-D, Zhou et al., 2015). In addition, ISC proliferation phenotypes were confirmed by multiple independent RNAi lines (Fig. S1E) and were also observed after tissue-specific CRISPR knockout of Mad, Med (Fig. S1Q). Together, these results show that inactivation of BMP signaling causes accumulation of stem cells and a multilayered epithelium, resembling previously described intestinal tumors induced by dysregulation of Notch and Sox21a (Patel *et al*, 2015; Zhai *et al*, 2015).

**Figure 1.**
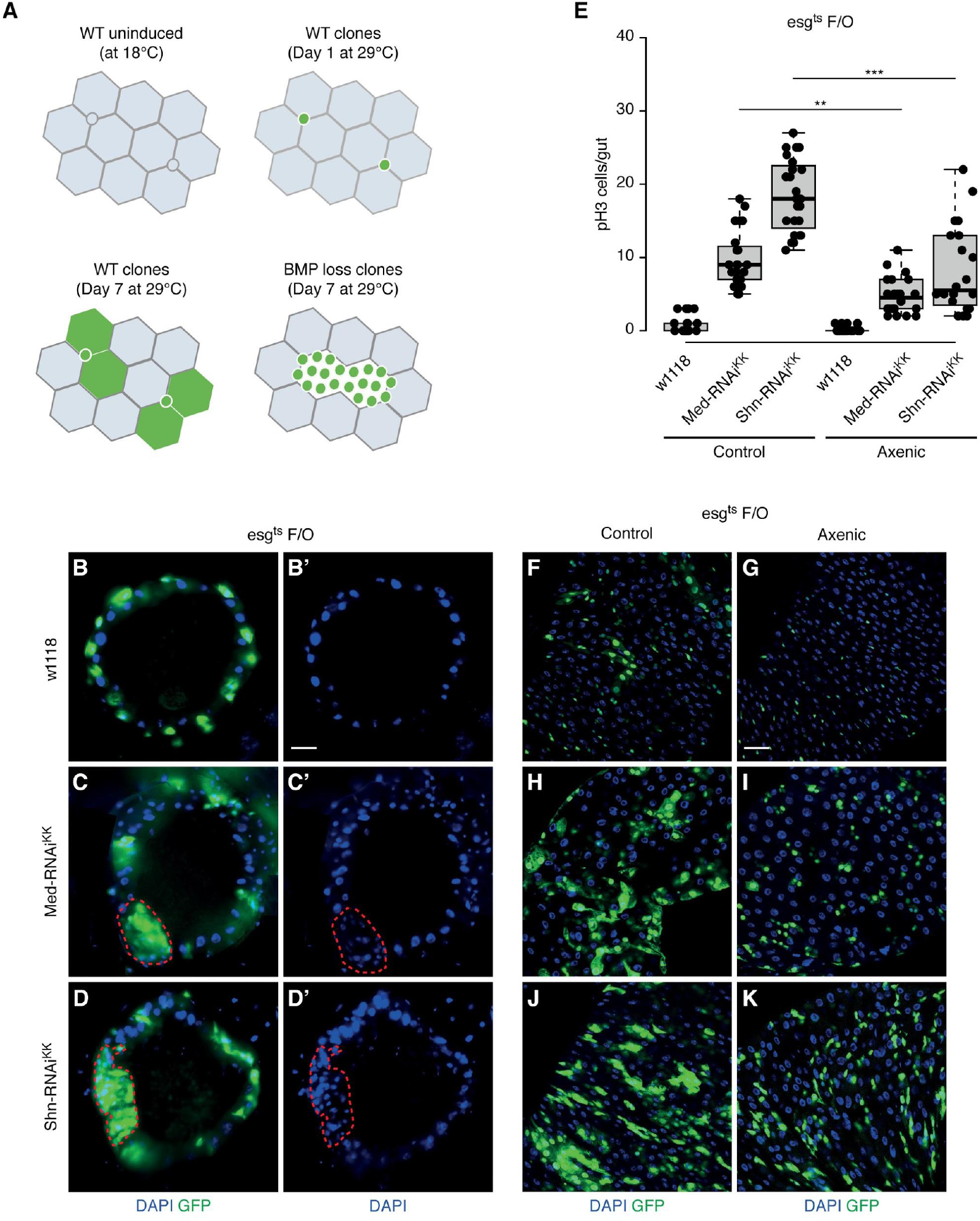
The intestinal microbiome promotes tumorigenesis. (A) The diagram displays the *esg*^*ts*^*F/O* system. (Upper left panel) the *F/O* system is inactivated by the temperature-sensitive repressor Gal80^ts^ at 18°C. (Upper right panel) At day 1, Gal80^ts^ is suppressed at 29°C and esg-Gal4 will drive the expression of UAS-GFP and UAS-flippase, in turn, activating the Act>CD2>Gal4 cassette. (Lower left panel) At day 7, the F/O system will express UAS-GFP and transgene of interest in the progenitor cells from stem cells and enteroblast due to the ubiquitous Act-Gal4 expression. (Lower right panel) At day 7, the F/O system activated Mad, Med or Shn RNAi transgene and GFP, causes accumulation of small nuclei GFP positive cells. (B-D) Representative images of cross sections of *esg*^*ts*^*F/O>w1118* (B), Med-RNAi (D) and Shn-RNAi (D) midgut after 10 days at 29°C. GFP (green): *Act-GFP*, DAPI (blue): DNA. Flip-out clones are indicated by a dashed red line. (E) Quantification of pH3-positive cells per adult midgut of the indicated genotypes in standard and antibiotics-treated fly food after 5 days at 29°C. (F-K) Representative images of the posterior midgut of *esg*^*ts*^*F/O>w1118* (F, G), *esg*^*ts*^*F/O>Med-RNAi*^*KK*^ (H, I) and *esg*^*ts*^*F/O>Shn-RNAi*^*KK*^ (J, K) flies raised in axenic condition, stained with pH3 antibody in red. *Act-GFP* is shown in GFP, nuclei are stained with DAPI (blue). * p<0.05; ** p<0.01; *** p<0.001. Scale bars: 30 μm (A-C, D-I, K-N)

Next, to gain insights into the contribution of the microbiota to intestinal tumor growth, we depleted the intestinal microbiome by raising flies under axenic condition or feeding antibiotics. We observed that Med or Shn RNAi animals raised under germ-free conditions developed fewer *esg*^+^ cell clusters and showed a significant decrease in stem cell mitosis (Fig. S1E-K). In addition, microbiota depletion reduces tumor related increase in stem cell proliferation in Shn RNAi flies (Fig. S1F-L). Consistently, knockouts of Med and Mad by tissue-specific CRISPR/Cas9 also showed a significant reduction in tumor growth upon antibiotic feeding as compared to control conditions (Fig. S1M-Q). We confirmed that antibiotic treatment significantly reduced the gut microbiome in control and Shn RNAi flies (Fig. S1R). Taken together, these data indicate that the depletion of the microbiome impacts stem cell proliferation and inhibits tumor growth.

### Intestinal tumors influence the microbiota

To assess whether the induction of intestinal tumors alters the microbiome, we first monitored changes in gut bacterial load. Interestingly, we found a significant increase in bacterial load upon Mad, Med and Shn RNAi as well as after tissue-specific Med knock-out (Fig. 2A and B), in addition, we confirmed an age-dependent increase in the Shn RNAi intestine (Fig. S2A).

**Figure 2.**
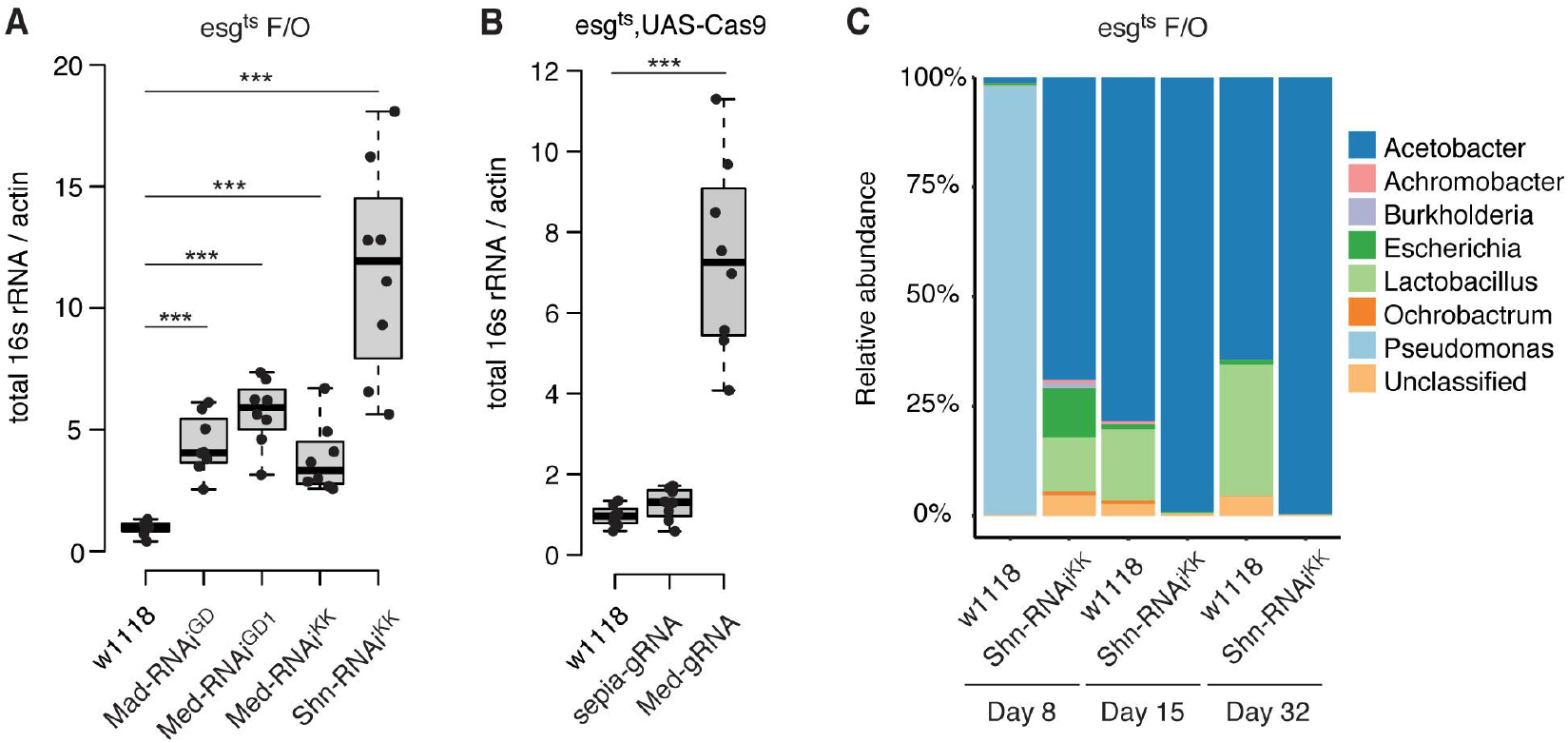
Intestinal tumors induce changes in the microbiome. (A) Bacterial levels in the intestines of *esg*^*ts*^*F/O>Mad-RNAi*^*GD*^, *esg*^*ts*^*F/O>Med-RNAi*^*GD-1*^, *esg*^*ts*^*F/O>Med-RNAi*^*KK*^, *esg*^*ts*^*F/O>Shn-RNAi*^*KK*^ and *esg*^*ts*^*F/O>w1118*, assayed by qPCR of 16s rRNA gene at 15 days at 29°C. (B) Bacterial levels in the intestines of *esg*^*ts*^;*UAS-Cas9 > Med-gRNA*, *esg*^*ts*^;*UAS-Cas9>sepia-gRNA* and *esg*^*ts*^; *UAS-Cas9>w1118*, assayed by qPCR of 16s rRNA gene at 18°C for 30 days after inducing mutagenesis. sepia-gRNA was used as a negative control. (C) The commensal composition of *esg*^*ts*^*F/O>Shn-RNAi*^*KK*^ and *esg*^*ts*^*F/O>w1118*midguts after 8, 15 and 32 days at 29°C. Bar charts show the top eight bacterial genera determined by 16S rRNA sequencing.

To further investigate changes in the microbial composition, we conducted 16S rRNA sequencing on Shn RNAi and control intestines. The experiments revealed an enrichment of *Acetobacteria* species concurrent with a significant reduction in overall microbial diversity in the intestine of tumor-bearing flies (Fig. 2C). Since the comparative analysis did not provide information about the extent changes in taxa abundance and changes in the overall microbiome (Vandeputte *et al*, 2017), we examined the load of various bacteria by CFU assays on selective mediums for *Acetobacteriaceae* and other bacteria (NR) (Ryu *et al*, 2008; Guo *et al*, 2014) as well as by taxa-specific 16S qPCR (Clark *et al*, 2015; Salazar *et al*, 2018). In these experiments, we observed intestinal dysbiosis in Dpp/BMP knock-down conditions (Mad/Med, Shn RNAi), as shown in Fig. S2B and C. A similar dysbiosis was observed in Notch-depleted (N RNAi) animals as an independent means to induce stem cell tumors (Fig. S2C). Furthermore, animals with intestinal specific Med knockouts showed a significant increase in *Acetobacteria* and other bacteria upon growth on nutrient rich medium (Fig. S2D). Together, these data demonstrate an overall increased bacterial load and a lower microbial diversity in the intestines of tumor-bearing flies.

Previous work showed that BMP signaling determines copper cell differentiation and its inactivation leads to decline of the acidic stomach-like copper cell region (CCR) in the middle midgut (Guo *et al*, 2013; Li *et al*, 2013, 2016). In addition, age-related CCR decline disrupts the distribution and diversity of gut commensals (Li *et al*, 2016). To exclude that tumor related microbial dysbiosis is caused by decline of the acidic stomach-like copper cell region (CCR), we assessed the function of the CCR by feeding control and tumor bearing flies with a diet containing a pH indicator. These experiments indicated that the tumor flies displaying dysbiotic phenotype at early time points (day 5 and day 8) but maintain the acidity in the CCR (Fig. S2A, E, F). In addition, the expression of CCR-specific vacuolar-type H+ ATPase, *Vha100-4* remained at a similar level in the tumor intestine compared to the control tissue (Fig. S2G). These results suggest that the tumor-induced dysbiosis is not caused by BMP-related decline in CCR function.

### Intestinal tumorigenesis impairs epithelial barrier functions

The complex rearrangements of cell-cell adhesion are required for shaping the intestine and for maintaining tissue integrity during development (Bryant & Mostov, 2008; Oh *et al*, 2012). The maintenance of the intestinal barrier is essential for preserving homeostasis and organismal health (Clark *et al*, 2015; Salazar *et al*, 2018). To test whether loss-of BMP-induced intestinal tumors disrupts barrier function by altering cell adhesion molecules, we monitored the expression of septate junction proteins Disc large (Dlg) and Fascillin III (Fas III) using immunostaining. We observed significant changes in the septate junction proteins (Dlg, FasIII) with protein loss and mislocalization (Fig. 3A-F, Fig. S3A and B). These results show that the septate junctions between tumor and intestinal cells are disrupted in the tissue.

**Figure 3.**
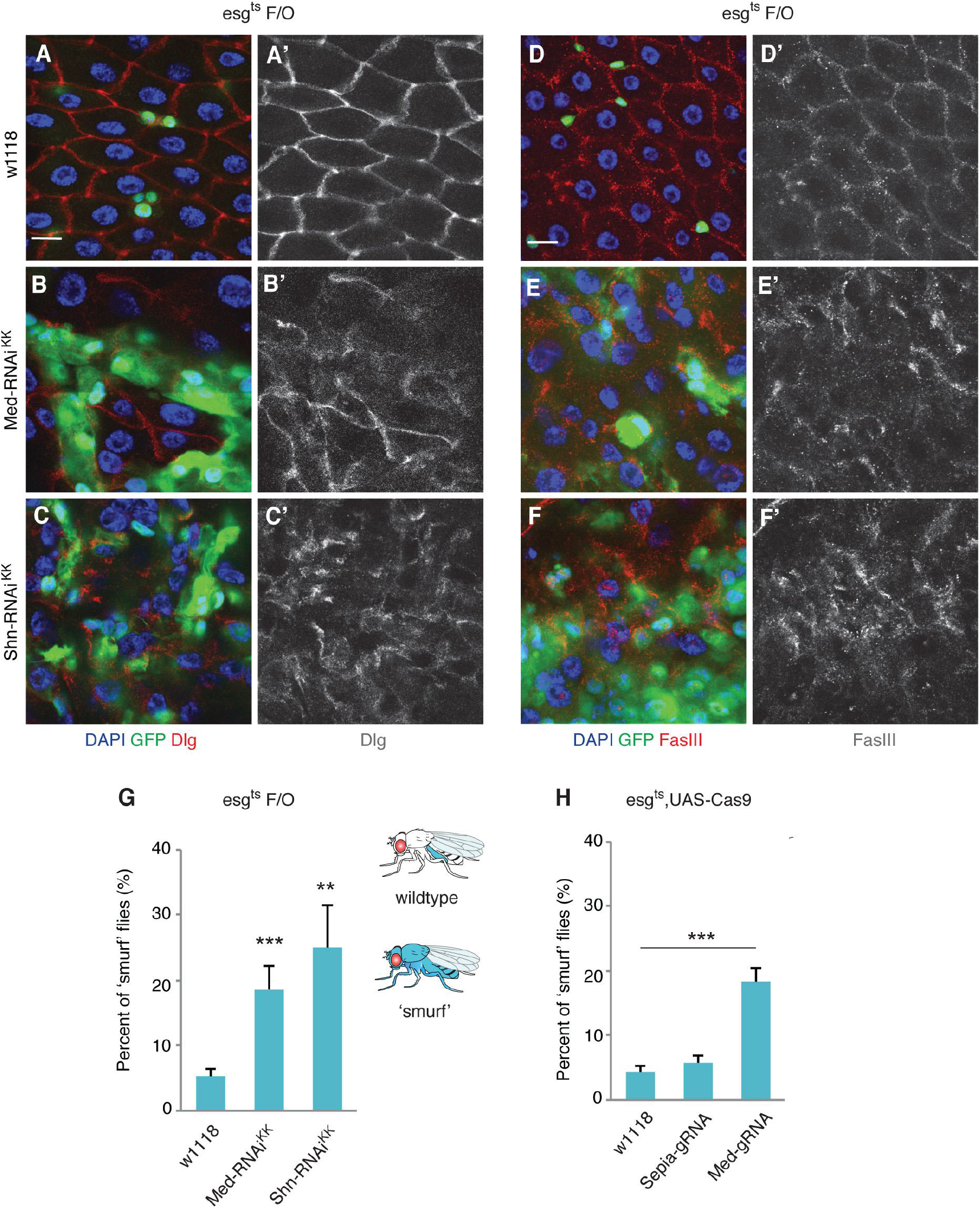
Tumor growth leads to disruption of epithelial barrier function. (A-C) Representative images of the posterior midgut of *esg*^*ts*^*F/O>w1118* (A), *esg*^*ts*^*F/O>Med-RNAi*^*KK*^ (B), *esg*^*ts*^*F/O>Shn-RNAi*^*KK*^ (C) flies at 29°C for 7 days, stained with anti-Disc large (Dlg) antibodies. (D-F) Representative images of the posterior midgut of *esg*^*ts*^*F/O>w1118* (D), *esg*^*ts*^*F/O>Med-RNAi*^*KK*^ (E), *esg*^*ts*^*F/O>Shn-RNAi*^*KK*^ (F) flies at 29°C for 7 days, stained with anti-Fascillin III (FasIII) antibodies. (G-H) ‘Smurf assay’ for gut integrity and barrier defects of indicated genotype. A high percentage of ‘smurf’ flies are shown in the group of *Med-RNAi*^*KK*^, *Shn-RNAi*^*KK*^ (G) or *Med-gRNA*(H) flies compared to the controls (*w1118* or *sepia-gRNA*). The “smurf” assay on RNAi or CRISPR experiment is monitored at 29°C for 25 days or at 18°C for 45 days after inducing mutagenesis. * p<0.05; ** p<0.01; *** p<0.001. Scale bars: 10 μm (A-F)

To further explore whether tumor-bearing flies show barrier defects, we fed flies with a blue dye to monitor intestinal epithelial integrity (also referred to as ‘smurf’ assay, (Rera *et al*, 2012). Flies with tumor frequently showed impaired intestinal barriers, with a significantly higher fraction of non-absorbable blue dye leakage to the fly compared to controls (Fig. 3G and H). These results suggest that tumor growth induces epithelial barrier defect. We also observed that Med and Shn depleted midguts were shorter when compared to control midguts (Fig. S3C-E, H). Similarly, Med CRISPR knockouts also showed a shorter gut phenotype (Fig. S3F, G, I).

Clark et al. (2015) reported that intestinal barrier loss leads to immune activation and increased tissue regeneration. To further test whether tumor growth impairs barrier function, we examined in more detail the expression of genes involved in immune activation (Peptidoglycan Recognition Protein SC2, PGRP-SC2; Diptericin, Dpt), reactive oxygen species (ROS) production (Duox), cytokine for STAT signaling activation (Upd3) and stress signaling (Kayak [Kay], the homolog of Fos) in the intestines of Shn RNAi flies. Quantitative RT-PCR from dissected intestine revealed that the JNK signaling pathway target Kay was highly induced in Mad/Med-RNAi or Shn-RNAi flies, as well as the cytokine Upd3 (Fig. S3J). We also observed a decrease in PGRP-SC2 expression and an increase in Dpt levels, indicative of immune activation in Mad/Med-RNAi or Shn-RNAi intestines (Fig. S3J and K). PGRP-SC2 is a known negative regulator of the Relish pathway and its suppression has been linked to commensal dysbiosis (Guo *et al*, 2014). We further found that the NADPH oxidase (Duox), responsible for producing microbial ROS in response to infection (Lee *et al*, 2015), showed a significantly decreased expression in the tumor intestine (Fig. S3K). Together, these results indicate that intestinal tumors lead to activation of stress and immune signaling, short gut length and impairs intestinal barrier function.

### JNK activation contributes to tumor growth

To understand the mechanisms promoting tumor growth and dysbiosis, we next assessed JNK signaling activity in Med or Shn depleted midguts. JNK was activated in the intestine after Med or Shn depletion, as measured by *Kay* and *puc* upregulation (Fig. S3J). Further experiments using a *puckered-lacZ* (*puc-lacZ*) reporter showed that JNK signaling was activated not only within the tumor but also in the nearby ECs (Fig. 4A-C). Moreover, CRISPR/Cas9 mediated Mad or Med intestinal knock-outs showed a similar upregulation of *puc-lacZ* signal (Fig. 4E-G). Consistently, we observed an increase of phospho-JNK levels in tumor intestine (Fig. S4A-C).

**Figure 4.**
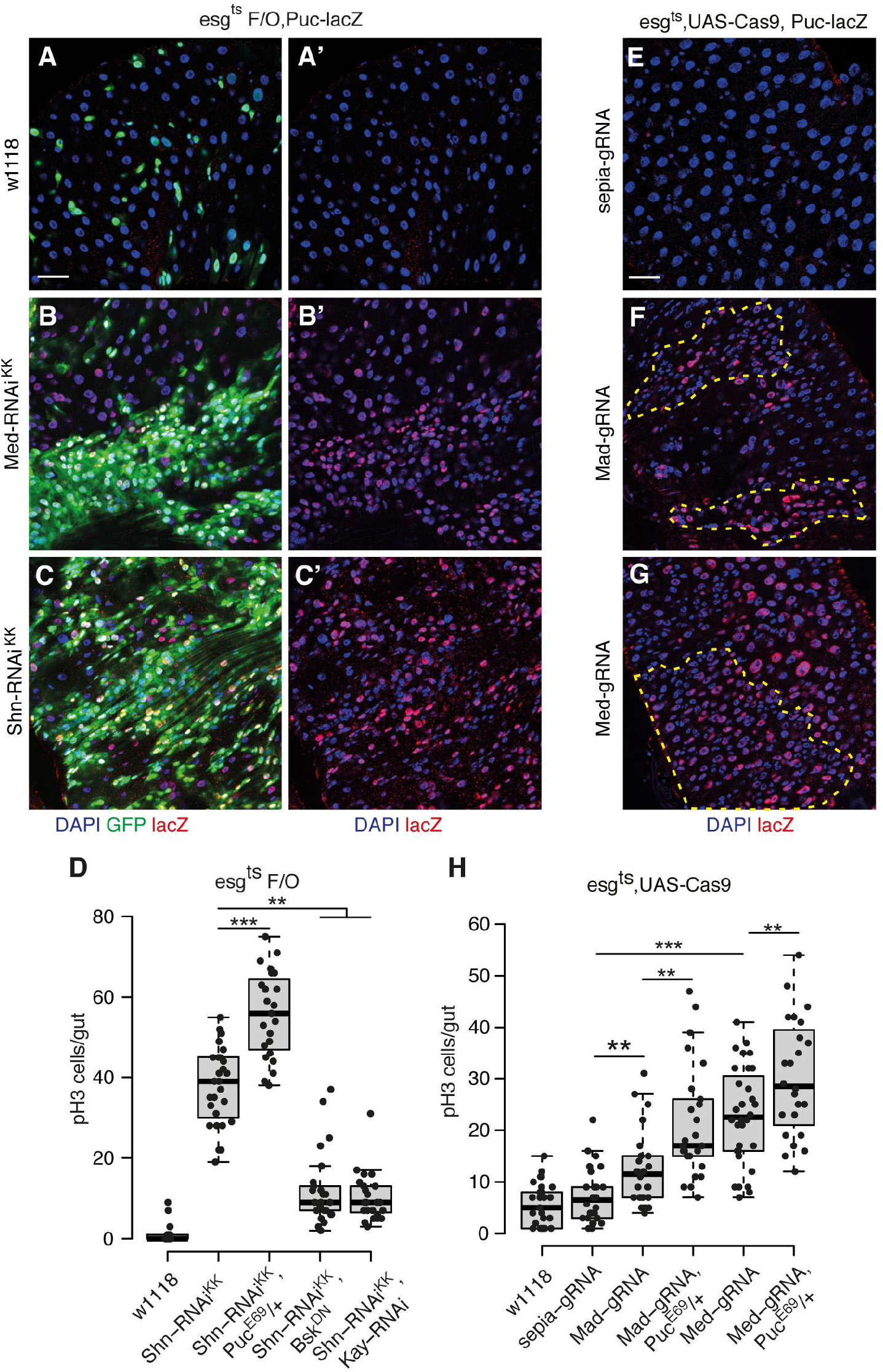
BMP-loss induced tumors activate JNK. (A-C) The *puc-lacZ* expression in the midgut of *esg*^*ts*^*F/O>w1118*(A), *esg*^*ts*^*F/O>Med-RNAi*^*KK*^ (B) and *esg*^*ts*^*F/O>Shn-RNAi*^*KK*^ (C) with anti-beta Galactosidase staining at 29°C for 7 days. (D) Quantification of pH3-positive cells per adult midgut of the indicated genotypes at 29°C for 7 days. (E-G) Immunofluorescence images of midguts of *esg*^*ts*^;*UAS-Cas9 > Mad-gRNA* (F) or *Med-gRNA* (G) or *sepia-gRNA* (E) with anti-beta Galactosidase staining at 18°C for 30 days after inducing mutagenesis. (H) Quantification of pH3-positive cells per adult midgut of the indicated genotypes at 18°C for 30 days. * p<0.05; ** p<0.01; *** p<0.001. Scale bars: 30 μm (A-C, E-G)

We next asked whether JNK signaling is required for BMP-dependent tumor growth. To this end, we either inhibited or activated JNK signaling by expressing a dominant negative form of *Basket* (*Bsk^DN^*), by *Kay RNAi*, or by the introduction of a heterozygous mutation of *Puckered* (*Puc^E69^/+*), a negative JNK pathway regulators (Martín-Blanco *et al*, 1998). We found that activating JNK signaling by removing one copy of Puckered significantly induced ISC division and tumor load in the Shn depleted intestine (Fig. 4D and Fig. S4D-H). Conversely, JNK suppression (*Bsk*^*DN*^ or *Kay-RNAi*) largely reduced ISC proliferation and tumor burden (Fig. 4D and Fig. S4D-H). In addition, JNK activation induced Mad or Med knock-out related tumor phenotype with elevated stem cell mitosis (Fig. 4H). These results suggest the activation of JNK contributes to intestinal tumorigenesis.

### JNK signaling contributes to intestinal barrier loss and dysbiosis

To test whether tumor-related JNK activation induces intestinal barrier dysfunction, we monitored the septate junction proteins and tissue integrity in the tumor bearing flies after modulation of JNK signaling. We found that JNK activation (*Puc*^*E69*^/*+*) enhanced the mislocalization and loss of septate junction proteins (Dlg, FasIII) in *esg F/O* driven Shn RNAi flies (Fig. 5A-C and Fig. S4I-K). In contrast, JNK suppression partially rescued the loss of septate junction protein phenotype in the tumor intestine (Fig. 5D). Shn RNAi flies with JNK activation displayed an even shorter midgut and this phenotype was completely suppressed by JNK inhibition (Bsk^DN^ or Kay-RNAi) (Fig. S4D-H, L). JNK suppression (*Bsk*^*DN*^ or *Kay-RNAi*) also extended the lifespan of flies with tumor (Fig. S4M and N), while JNK activation enhanced the mortality of tumor-bearing flies (Fig. S4M and N).

**Figure 5.**
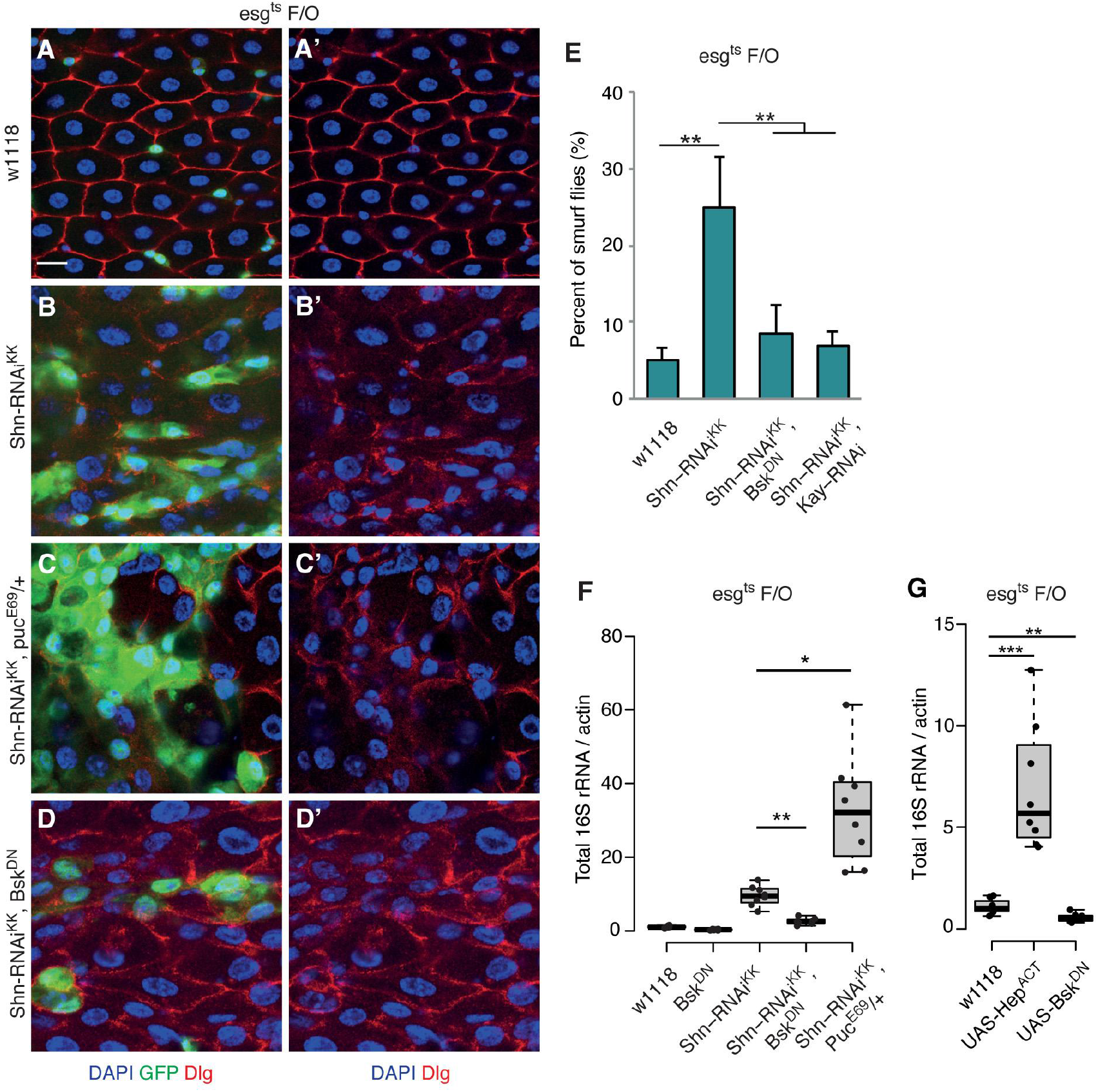
Tumor-induced JNK activation disrupts epithelial function and causes dysbiosis. (A-D) Representative images of the posterior midgut of *esg*^*ts*^*F/O>w1118*(A), *esg*^*ts*^*F/O>Shn-RNAi* (B), *esg*^*ts*^*F/O>Shn-RNAi*; *Puc*^*E69*^/+ (C), *esg*^*ts*^*F/O>Shn-RNAi*; *UAS-Bsk*^*DN*^ (D) flies stained with Dlg antibody. (E) ‘Smurf assay’ for gut integrity and barrier defects of indicated genotype at 29°C for 25 days. (F) The bacterial level in the intestine of *esg*^*ts*^*F/O>UAS-Bsk*^*DN*^, *esg*^*ts*^*F/O>Shn-RNAi*^*KK*^, *esg*^*ts*^*F/O>Shn-RNAi*^*KK*^;*UAS-Bsk*^*DN*^, *esg*^*ts*^*F/O>Shn-RNAi*^*KK*^, *Kay-RNAi* and *esg*^*ts*^*F/O>w1118*, assayed by qPCR of 16s rRNA gene at 29°C for 15 days. (G) Bacterial load in the intestine of *esg*^*ts*^*F/O>UAS-Hep*^*ACT*^, *esg*^*ts*^*F/O>UAS-Bsk*^*DN*^ genotypes, assayed by qPCR of 16s rRNA at 29°C for 12 days. Significant differences are indicated with asterisks (* p<0.05; ** p<0.01; *** p<0.001). Scale bars: 10 μm (A-D).

To explore how JNK activation contributes to increased mortality in tumor-bearing flies, we fed flies with blue dye to monitor intestinal epithelial integrity. In tumor-bearing flies, we observed a significant breakdown of tissue integrity and decreasing JNK activity resulted in a reduced proportion of ‘smurf’ phenotypes (Fig. 5E and Fig. S5O). In Sox21a tumors, JNK mediated Mmp2 expression can contribute to enterocyte detachment (Zhai *et al*, 2015). To ask whether Mmp2 is required for tumor growth, we co-expressed Shn RNAi and Mmp2 RNAi using the *esgts F/O* system. We observed a significant reduction in stem cell proliferation and tumor growth upon Mmp2 depletion (Fig. S5P). These results suggest that JNK is activated to trigger intestinal barrier dysfunction and accelerates tumor growth, probably through regulating its downstream effector Mmp2.

As the intrinsic JNK activity contributes to BMP-loss induced tumorigenesis and tissue failure, we hypothesized that JNK is also involved in tumor related microbiota dysbiosis. We next examined the overall bacterial load in the tumor tissue with aberrant JNK activity by 16S rRNA qRT-PCR. Blocking JNK activity by expressing Bsk^DN^ suppressed the increase in microbial load in the tumor bearing flies (Fig. 5F). In contrast, JNK activation (*Puc*^*E69*^/*+*) enhanced the tumor-related dysbiosis (Fig. 5F). Furthermore, the number of CFUs of two main microbiota phylotypes (*Acetobacter*, *Lactobacilli*) increased significantly in tumor bearing flies. In contrast, JNK suppression largely reduced the increased *Acetobacter*, *Lactobacilli* and other bacteria populations (Fig. S5Q). Indeed, the level of intestinal JNK activity is associated with the changes of microbiota (Fig. 5G). Hence, these results indicate that high intestinal JNK activity is linked to loss of host-microbe homeostasis.

### Intestinal tumor regression restores epithelial function and dysbiotic phenotype

To explore the temporal link between intestinal tumorigenesis, changes in immune responses and a dysbiotic microbiome, we designed a time-course experiment using the temperature controlled *esg*^*ts*^*F/O* system. First, we initiated tumor growth by depletion of Med or Shn for 7 days (‘T_7_’), and subsequently divided the population of flies into two groups that either continuously expressed (‘T_14_’) or stopped to express the Shn or Med RNAi transgenes (‘T_7+7_’) (Fig. 6A). We then monitored the expression of genes involved in immune activation (PGRP-SC1a, -b, PGRP-SC2, Dpt), stress signaling (Kay, Puc, Mmp1, Mmp2), ROS production (*Duox*), internal bacterial load, tumor growth and tissue morphology.

**Figure 6.**
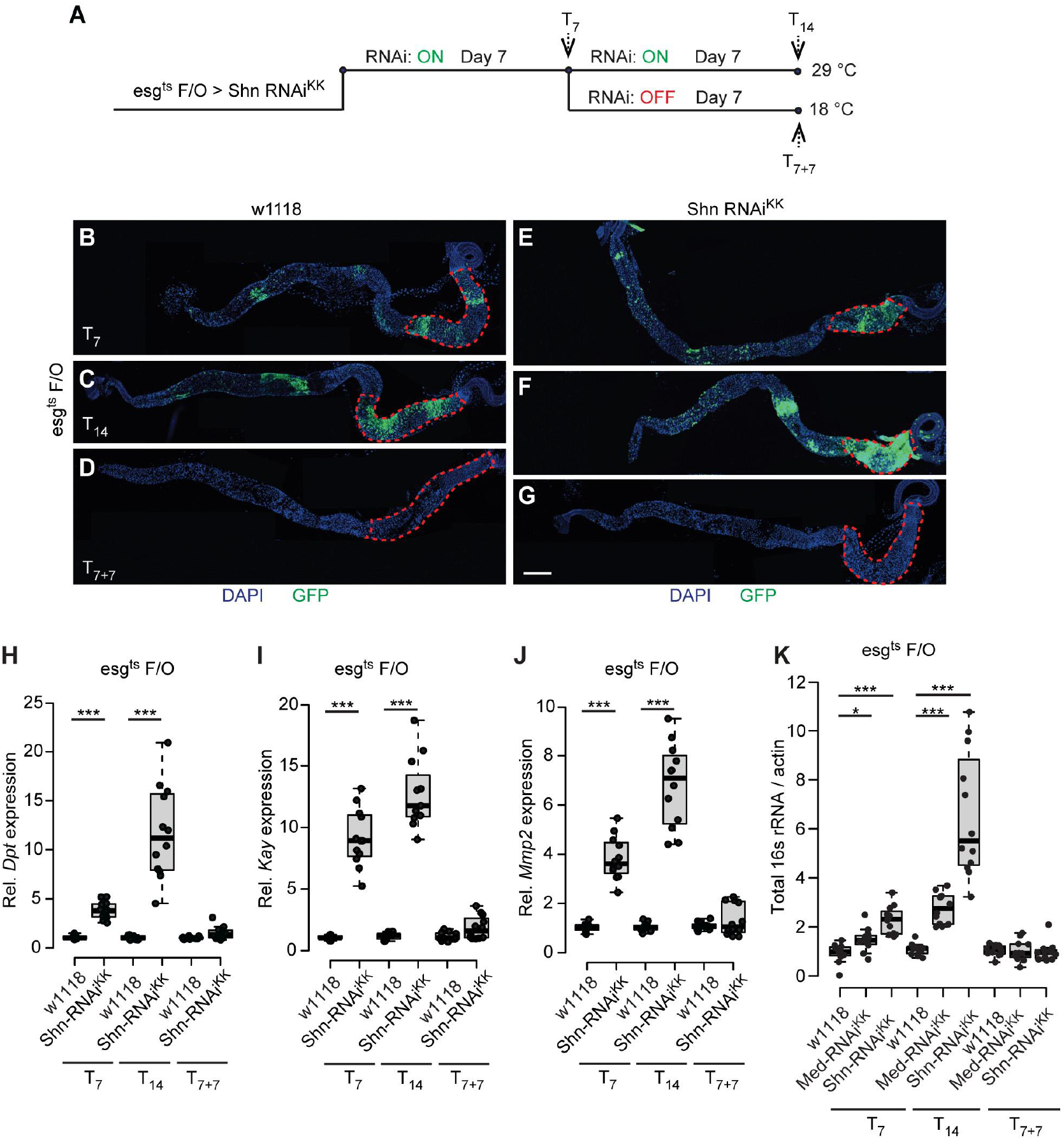
Regression of intestinal tumor restores epithelial function and reduces the dysbiosis. (A) The system to express transgenes in stem cell and its progenitors to initiate intestinal tumor and subsequently remove the tumor in a temperature control manner. (B-G) Representative images of midguts of *esg*^*ts*^*F/O>w1118*, *esg*^*ts*^*F/O>Shn-RNAi*^*KK*^ flies ‘T_7_’ (B, E) or ‘T_14_’ (C, F) or ‘T_7+7_’ (F, G). Conditions: Tumor start (T_7_): at 29°C for 7 days; Tumor progression (T_14_): at 29°C for 14 days; Tumor recovery (T_7+7_): at 29°C for 7 days then shifted to 18°C for another 7 days. *Act-GFP*is shown in GFP, nuclei are stained with DAPI (blue), please see the experimental details in Figure S4A. (H-J) RT-qPCR analysis of the intestine shows that the expression levels of Relish (*Dpt*-H) and JNK (*Kay*-I, *Mmp2*-J) pathway component genes are altered in *esg^ts^F/O>Shn-RNAi* midgut in the conditions of tumor ‘start’ (RNAi on at 29°C for 7 days), ‘progression’ (at 29°C for 14 days) and ‘recovery’ groups (at 29°C for 7 days, then shift to 18°C that switch off RNAi for 7 days). * p<0.05; ** p<0.01; *** p<0.001. (K) Bacterial levels in the intestine of *esg^ts^F/O>Shn-RNAi* and*esg*^*ts*^*F/O>w1118*, assayed by qPCR of 16s rRNA in tumor ‘start’, ‘progression’ and ‘recovery’ conditions. Scale bars: 250 μm (B-G)

These experiments showed that the continuous Shn RNAi caused an enhanced tumor growth with more severe tissue morphology defects (T_14_, Fig. 6F) compared to T_7_ (Fig. 6E). This tumor phenotype can be reversed in the ‘:T_7+7_’ condition by switching off RNAi expression and thereby restoring BMP signaling (Fig. 6G). In controls, no tumor phenotypes were observed at T_7_, T_14_ and T_7+7_ (Fig. 6B-D). Concurrently, we also observed a reversal of the increased stem cell proliferation and ‘shorter gut’ phenotype in the ‘T_7+7_’ group (Fig. S5A and B).

At the molecular level, we found that immune activation in the tumor tissue was restored to the control levels in the tumor ‘T_7+7_’ condition, as indicated by the expression of PGRP-SC1a, PGRP-SC1b, PGRP-SC2 and Dpt (Fig. 6H and Fig. S5C-E). Furthermore, the expression of JNK signaling target genes (Kay, Puc, Mmp1, Mmp2) was also reduced to control levels in the ‘T_7+7_’ condition (Fig. 6I-J and Fig. S5F-G). Duox expression was also significantly lower in the ‘T_7_’ and ‘T_14_’ groups but returned to normal levels in the ‘T_7+7_’ group (Fig. S5H). An increase in intestinal bacterial load was observed in ‘T_7_’ and ‘T_14_’ condition, whereas the bacterial load in the ‘T_7+7_’ conditions returned to a normal level as compared to the control group as well as the ISC proliferation (Fig. 6K). To further confirm the changes of *Alphaproteobacteria* and *Bacillus* bacteria families in the tumor ‘recovery’ group, we used specific primers as described previously (Sebald *et al*, 2016) and found that both *Acetobacteraceae* and *Lactobacillus plantarum* levels are increased in the ‘T_7_’ and ‘T_14_’ and returned to a normal level in the ‘:T_7+7_’ condition (Fig. S5I-J). Thus, our data indicate that the altered gut length, increased ISC proliferation and microbiota dysbiosis phenotypes are associated with tumor progression and can be restored when tumor growth is suppressed.

### Removal of microbiota restores intestinal function in tumor flies

To determine whether preventing tumor related dysbiosis can influence intestinal barrier dysfunction and animal health, we treated tumor animals with antibiotics to eliminate bacterial growth at 2 days prior to tumor initiation. The antibiotic treatment caused a significant reduction (p<0.05) in genes involved in stress signaling (Kay) and regenerative response (Upd3) in the tumor tissue (Fig. 7A and B). The expression of the antimicrobial peptide (Dpt) was also reduced in the gut of tumor flies upon antibiotics feeding (Fig. 7C). In addition, antibiotic treated tumor animal showed a reduced intestinal barrier failure (Fig. 7D). The antibiotic feeding also significantly increased the lifespan of Shn-RNAi flies (Fig. 7E), indicating tumor-related microbiota is detrimental to barrier-loss mediated mortality.

**Figure 7.**
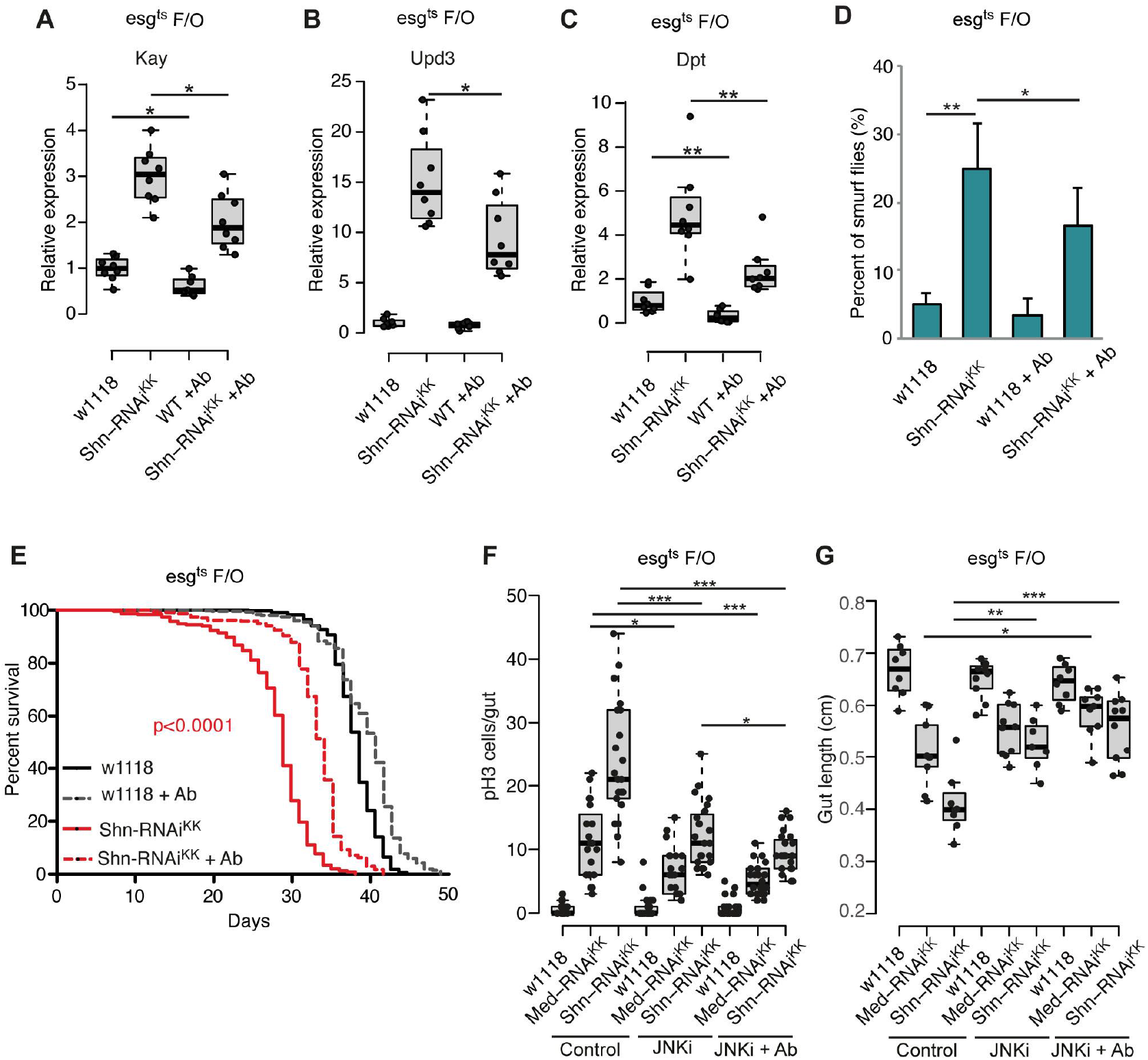
Depletion of the microbiome or inhibition of JNK signaling restores intestinal function and organismal health. (A-C) RT-qPCR analysis of the intestine shows that the expression levels of JNK (*Kay*, A), regenerative cytokine (Upd3, B) and Relish (*Dpt*, C) pathway component genes are reduced in *esg*^*ts*^*F/O> Shn-RNAi*^*KK*^ midgut upon antibiotic feeding at 29°C for 7 days. (D) Smurf assay for gut permeability of indicated genotype at 29°C for 25 days. Antibiotics feeding reduces Shn-RNAi induced Smurf phenotype. (E) The lifespan of Shn-RNAi tumor flies compared to control flies on normal or antibiotic food. No antibiotics: w1118 n=341, Shn-RNAi^*KK*^ n=234; with antibiotics: w1118 n=401, Shn-RNAi^*KK*^n=239. (F) Quantification of pH3-positive cells per adult midgut of the indicated genotypes after 8 days RNAi induction in control, JNK inhibitor (SP600125) feeding and a combination of JNK inhibitor and a cocktail of antibiotics. (G) Quantification of the length of intestine of *esg*^*ts*^*F/O>Shn-RNAi*^*KK*^, *esg*^*ts*^*F/O>Med-RNAi*^*KK*^ and *esg*^*ts*^*F/O>w1118* after 8 days RNAi induction in control, JNK inhibitor (SP600125) feeding and a combination of JNK inhibitor and a cocktail of antibiotics. * p<0.05; ** p<0.01; *** p<0.001.

To better understand the relationship between intrinsic JNK activation and extrinsic microbiota on intestinal tumor growth, we utilized antibiotic feeding in combination with the JNK inhibitor SP600125. Treatment with a JNK inhibitor alone showed a reduced level of stem cell division and tumor burden and led to a recovery on the gut length of tumor bearing flies (Fig. 7F and G). Interestingly, a combination of antibiotic treatment and JNK inhibitor feeding showed an additive effect on the suppression of stem cell activity and tumor growth (Fig. 7F). In addition, the combinatorial treatment showed a better recovery of the gut length phenotype in the tumor flies (Fig. 7G). Hence, these results indicate intestinal microbiota and intrinsic JNK activity are both required for tumor growth, whereas a combinatorial targeting shows a beneficial effect on tumor inhibition.

## DISCUSSION

Recent studies have investigated tumors that arise by loss of differentiation, accumulation of differentiating stem cell or stem cell hyperproliferation in adult *Drosophila* intestine (Guo *et al*, 2013; Patel *et al*, 2015; Ragab *et al*, 2011; Suijkerbuijk *et al*, 2016; Zhai *et al*, 2015; Cordero *et al*, 2012; Parvy *et al*, 2018; Siudeja *et al*, 2015; Kwon *et al*, 2015). Loss-of BMP signaling through mutation in SMAD4 or BMPR1A is a known cause of gastrointestinal cancers in man (Hardwick *et al*, 2008). Similarly, inactivation of BMP pathway components also leads to intestinal tumor phenotypes in *Drosophila* (Guo *et al*, 2013). In this study, we used an inducible intestinal tumor model by independent RNAi constructs or CRISPR/Cas9 mutagenesis on BMP signaling to investigate host-microbe interactions. Our experiments demonstrate that changes in the microbiome are associated with intestinal tumor formation and highlight an important role of the epithelial barrier in maintaining host-microbe homeostasis. We provide several lines of evidence that intestinal tumorigenesis results in severe barrier defects, which, in turn, cause dysbiosis. The dysbiotic microbiome and tumor cells form a feedback loop to further fuel tumor growth. Mechanistically, we show that tumor growth alters the expression of genes involved in the control of microbiota in the intestine, including PGRP-SC2 and Duox. Furthermore, tumorigenesis causes loss of septate junction proteins, which were previously shown to be required for strengthening the epithelium structure and form a barrier to solute diffusion through the intercellular space (Hijazi *et al*, 2009). In addition, tumor growth alters gut morphology and give rise to a shrunken tissue. Finally, a growing tumor tissue leads to epithelium breakdown and increased intestinal permeability.

Functional and structural disruptions of the intestinal barrier trigger alterations in both bacterial load and composition and thereby induce commensal dysbiosis. We show that intestinal barrier failure is associated with an increase in JNK activity. Our data further support a model whereby tumor-related JNK activation is responsible for intestinal barrier dysfunction, leading to commensal dysbiosis as well as immune activation and a subsequently shorter lifespan. Inhibition of tumor growth or JNK inhibition reverses these phenotypes. Moreover, microbial removal restores intestinal barrier function and delays tumor growth, which can extend the lifespan of tumor-bearing animals.

### Self-enforcement between tumor and dysbiotic microbiome

In *Drosophila*, intestinal commensals have been recognized as important factors that influence gut immunity and homeostasis (Buchon *et al*, 2013b; Lee & Brey, 2013). For instance, microbiota mediated ERK signaling in ECs is required for epithelial response to restrict intestinal viral infections (Sansone *et al*, 2015). Our data show that tumor growth causes the reduction of microbial diversity. Previous studies indicated that reduced microbial diversity may increase the risk of susceptibility to infection and stress. In humans, a significantly lower microbiota diversity has been reported in colorectal cancer (Wu *et al*, 2013; Chen *et al*, 2012). Furthermore, several bacterial species such as *Streptococcus bovis*, *Helicobacter pylori*, *Bacteroides fragilis*, *Enterococcus faecalis*, *Clostridium septicum*, *Fusobacterium spp.* and *Escherichia coli* have been associated with colorectal carcinogenesis (Gagnière *et al*, 2016), although the causal link often remains unclear. Our data also demonstrate that microbial clearance reduces proliferative and inflammatory signals in tumor intestine with a decreased stem cell activity, indicating the dysbiotic microbiome create a pro-inflammatory environment that promotes tumor growth.

### Tumor growth causes intestinal dysfunction and dysbiosis

In response to infection, intestinal epithelium undergoes severe morphological changes, while retaining the ability to recover within a few hours (Lee *et al*, 2016). Barrier dysfunction has been correlated with impaired insulin and immune signaling (Rera *et al*, 2012). Our experiments show that tumor growth impairs the intestinal renewal capacity and barrier function by inducing gut integrity loss, immune activation, reduced Duox expression and in turn causes intestinal dysbiosis. This is consistent with previous results showing that intestinal barrier failure after loss of a septate junction protein Snakeskin results in altered gut morphology, dysbiosis and reduced Duox level (Salazar *et al*, 2018). Duox act as an important defence mechanism in ECs against enteric infection and its expression is increased in the intestine during infection and ageing (Lee *et al*, 2018; Guo *et al*, 2014), suggesting that Duox expression is induced during normal intestinal regeneration under infection or ageing, while its expression is dampened in the tumor bearing intestine with massive loss of ECs and impaired EC renewal capacity. A recent study demonstrated lactic acid from *L. plantarum* activates Nox for ROS production that promotes intestinal damage and dysplasia (Iatsenko *et al*, 2018). Increased ROS level is also detected in the vicinity of Sox21a tumor (Zhai *et al*, 2015). However, molecular evidence on how the ROS level is regulated during tumorigenesis is still lacking and further studies are needed to investigate whether an increased microbiota also trigger intestinal ROS level through Nox for tumor progression.

Previous studies demonstrate that BMP signaling is required for CCR function and ageing related decline in CCR has been linked to microbial dysbiosis (Li *et al*, 2013, 2016). Our results indicate BMP loss tumor related intestinal dysbiosis is independent of CCR decline. The gastric stem cells in the CCR are largely quiescent only can be induced to regenerate the gastric epithelium upon challenge (Strand & Micchelli, 2011). We therefore speculate the CCR decline related microbiota changes occurs only at regenerative or ageing condition, whereas the tumor growth does not alter the CCR identity at a young age.

### JNK activation contributes to tumorigenesis and epithelial barrier failure

In this study, we showed the association of Mmp1 and Mmp2 expression with tumor progression and reducing Mmp2 expression dampens the tumor growth. Indeed, the expression of JNK downstream effectors Mmp1 and Mmp2 were induced in the EBs of *Sox21a* tumor. EB specific Mmp2 depletion in *Sox21a* flies reduced tumor burden and growth towards lumen (Zhai *et al*, 2015). The matrix metalloproteinases (MMPs) are involved in extracellular matrix (ECM) component cleavage (Kessenbrock *et al*, 2010). We therefore hypothesize that the JNK mediated MMP expression may be involved in extracellular matrix degradation and facilitates tumor growth and/or induces cell death to adjacent ECs. Therefore, tumor growth requires JNK related MMPs expression. In mammals, JNK activity affects cell proliferation, differentiation, survival and migration in a context-dependent manner and these events have also been linked to tumorigenesis (Bubici & Papa, 2014). Our data also indicate the increased JNK activity causes intestinal barrier failure. Therefore, dampening JNK signaling might be an effective strategy to prevent loss of barrier function, microbial dysbiosis and tumor progression.

### Conclusions

Our study underlines the importance of a tight regulation of JNK signaling for epithelial barrier function in order to maintain host-microbe homeostasis. Intestinal tumors activate JNK signaling to fuel tumor growth and barrier dysfunction. The impaired intestinal barrier disrupts host-microbe homeostasis and leads to dysbiosis, which initiate a proliferative and/or inflammatory response to further promote tumorigenesis. Interestingly, we found that JNK inhibition and/or microbiota depletion restores epithelial barrier function and reduces tumorigenesis. Therefore, combinatorial targeting of a tumor-induced microbial dysbiosis and stress signaling pathways in tumor cells could delay tumor growth and prevent intestinal organ dysfunction.

## MATERIALS AND METHODS

### *Drosophila* culture and medium

Animals were reared at either 18°C or 29°C with a 12 hours light cycle with 60% humidity. 1 liter standard fly medium contains: 44g sugar sirup, 80g malt, 80g corn flour premium G750, 10g soy flour, 18g yeast, 2.4g methly-4-hydroxybenzoate, 6.6 ml propionicacid, 0.66 ml phosphoricacid and 8g agar) as also previously described (Zhou *et al*, 2017). Antibiotic treatment was conducted by adding 150 μg/ml carbenicillin, 150 μg/ml metronidazole and 75 μg/ml tetracyclin to standard fly food. The antibiotic cocktail was added to the food at a suitable temperature to prevent heat inactivation. Animals in control or experimental groups are transferred to fresh food every two days to prevent fungal infection.

### Fly stocks

*esg*^*ts*^*F/O (Jiang et al, 2009)* (*esg-Gal4,UAS-GFP; Act>CD2>Gal4, TubGal80*), *MyoIA*^*ts*^, *MyoIA-lacZ (Jiang et al, 2011)*, UAS-Hep^ACT^, Puc^E69^ (Jiang *et al*, 2009), UAS-Notch-RNAi (Patel *et al*, 2015), *esg*^*ts*^ and Dl-lacZ (kind gift of Bruce Edgar), UAS-Upd2 (Zhai *et al*, 2015), UAS-Upd2-RNAi (Zhai *et al*, 2015), UAS-Kay-RNAi (Uhlirova & Bohmann, 2006), *UAS-Mmp2-RNAi* (Uhlirova & Bohmann, 2006) (kind gift of Mirka Uhlirova), (BLN6397), UAS-Mad-RNAi^Trip^ (BLN31315), UAS-Med-RNAi^KK^ (KK108412, VDRC), UAS-Shn-RNAi^KK^ (KK101278, VDRC), UAS-Mad-RNAi^KK^ (KK110517, VDRC), UAS-Mad-RNAi^GD^ (GD12635, VDRC), Med-RNAi^GD-1^ (GD19688, VDRC) Mad-RNAi^GD-2^ (GD19689, VDRC), UAS-Bsk^DN (Jiang *et al*, 2009)^ (BLN6409), UAS-Cas9 (Port *et al*, 2014) and guide-RNA transgenic strains (this study): *sepia-gRNA*, *Mad-gRNA*, *Med-gRNA*. Nubbin-Gal4 driven Mad-RNAi^Trip^, Med-RNAi^KK^ and Shn-RNAi^KK^ were previously tested to cause Dpp/BMP loss of function phenotype with small wings and vein formation loss (Zhou *et al*, 2015). Shn-RNAi results in stronger wing phenotype than Med or Mad RNAi (Zhou *et al*, 2015). The KK RNAi lines used in this study were tested negative by PCR for a *tiptop* off-target phenotype caused by an additional insertion at the 40D locus (data not shown) (Vissers *et al*, 2016).

### Axenic condition experiments

To generate germ-free flies, embryos were collected on grape juice plates. Briefly, embryos less than 12 hrs old were washed with 1X PBS and then rinsed in 70% ethanol. After that, embryos were bleached with 3% solution of sodium hypochlorite for 2-3 mins and washed with sterile water three times. Axenic embryos were then transferred to autoclaved medium in a sterile hood and maintained at 18°C. Subsequent generations were maintained in parallel to their conventionally-reared counterparts by transferring adults to new sterile tubes in a separate sterile box at 29°C for RNAi induction. Axenic conditions were confirmed by 16S rRNA bacterial qPCR as described previously (Buchon *et al*, 2009; Guo *et al*, 2014).

### *Drosophila* lifespan assay

One bottle containing 20 virgin females (*esg*^*ts*^*F/O*, *MyoIA*^*ts*^ drivers) and 5-7 males (RNAi transgenes or controls) were used to make a cross. Progenies were collected in 3 days after eclosion. Around 80 females plus 20 males with indicated genotype were transferred to a cage and incubated at 29°C to induce RNAi effect. At least 3 biological replicates for each genotype were prepared for each experiment. The food was changed every 2-3 days to prevent fly drown into food or fungal growth. Only the dead female flies were counted during ageing and the data were analyzed using PRISM 6 statistical software. The same cocktail of antibiotics as described above was used to generate antibiotic containing food for the life span assay on bacteria-depleted environment. The life span assay has been performed 3 times and only one experiment is shown in the related figure.

### Generation of sgRNAs transgenic stocks

Transgenic sgRNA stocks (Mad-gRNA-HD_CFD00651 and Med-gRNA-HD_CFD00386) were generated as part of the Heidelberg CRISPR fly library. Target sites were identified using E-CRISP (a web application to design gRNA sequences) (Heigwer *et al*, 2014) and pairs of sgRNAs, targeting the 5’ portion of the coding sequence, were cloned into pCFD6, using methods described previously (Port & Bullock, 2016). Plasmids were integrated at attP40 on the second chromosome using PhiC31 mediated integration and confirmed by Sanger sequencing. The gRNAs sequences are:

Mad-sgRNA1-TGTGTTCGATTCCACATCGT
Mad-sgRNA2-TGGCGAGGAACGAGTACTGG
Med-sgRNA1-GTGGGGGCGTTCGAGGGCGG
Med-sgRNA2-GTCCGGCCTGAGTCTGCAGT

### Tissue specific CRISPR/Cas9 mediated mutagenesis

The expression of Cas9 is temporally controlled by *UAS/Gal4/Gal80*^*ts*^ system in the intestinal stem cell compartments (*escargot-Gal4; Tubulin-Gal80*^*ts*^; *UAS-Cas9* and in short: *esg*^*ts*^, *UAS-Cas9*). The Mad or Med gRNAs male flies were crossed with the driver *esg*^*ts*^>*UAS-Cas9* at 18°C to prevent the mis-expression of Cas9 during the fly development. Their progeny with the right genotype (*esg*^*ts*^, *UAS-Cas9>UAS-t∷gRNA-Mad*^*2x*^ *or UAS-t∷gRNA-Med*^*2x*^ were collected and shifted to 29°C to activate Cas9 expression for 5 days. Afterward, the flies were maintained at 18°C until sacrifice for the experiments. Note that *esgts* driven Cas9 and sgRNAs is expressed only for 5 days to induce mutations and then animals are shifted to 18°C. No GFP expression is observed as *esg-GFP* is repressed by tubGal80 at 18°C. In general, we observed a higher phenotypic variability in CRISPR as compared to RNAi experiments which are likely caused by genetic mosaics of edited cells as previously described (Port and Bullock, 2016). To determine the CRISPR on-target efficiency, genomic DNA was extracted from 20 dissected intestine of CRISPR-Cas9 edited flies (genotype: esgts, UAS-Cas9>Med-gRNA) and the genomic locus surrounding the sgRNA target sites was amplified by PCR. PCR amplicons were subjected to Sanger sequencing and the resulting chromatographs were analyzed by Inference of CRISPR Edits (ICE) analysis (https://ice.synthego.com).

### Epithelial barrier assay and pH measurements

The barrier defect assay (‘smurf assay’) was conducted as described previously (Rera *et al*, 2012). Briefly, more than 60 flies were maintained on standard food until the day of ‘smurf assay’. Blue dye food was prepared by adding 2.5% (weight/volume) bromophenol blue (Sigma) to normal food, which was also used as pH indicator to assay the acidity of copper cell region as previously described in (Li *et al*, 2016). Flies were fed for 12 hrs and then monitored for the proportion of animals with a phenotype as an indicator for epithelial barrier dysfunction. To check for pH, intestines containing pH indicator were dissected in PBS. The head and posterior part were maintained intact to avoid the dye leakage. To avoid color change due to tissue incubation in PBS, images were taken immediately after dissection.

### Midgut RT-qPCR

RNA was extracted from 10 midguts (females) using RNeasy (Qiagen). One microgram of RNA was reverse transcribed using the RevertAid H Minus First Strand cDNA Synthesis Kit (Fermentas). RT-qPCR was performed in a 384-well format using the Universal Probe Library or SyBr green qPCR master-mix on a LightCycler 480 (Roche). qPCR primer sequences used in this study are listed in Table S2. At least 3 biological replicates (n≥3) and 2 independent experiments were performed. All results are presented as the means ±SD of the biological replicates. RP49 was used as a normalization control as previously described (Zhou et al, 2017).

### JNK inhibitor and antibiotic treatment

Flies were fed with 1 mg/ml JNK inhibitor SP600125 in 5% sucrose solution. For the combination of JNK inhibitor and antibiotics treatment, the same concentration of SP600125 was used plus an antibiotic cocktail (150 μg/ml ampicillin, 150 μg/ml metronidazole and 75 μg/ml tetracyclin) solved in 5% sucrose solution. Flies were fed on a round filter paper disc with SP600125, antibiotic cocktail or a combined solution and transferred to a freshly prepared vial every day to prevent evaporation effects.

### Intestinal microbiota assays by 16S rRNA qPCR and CFUs

The outside surface of animals was sterilized with 100% ethanol for 1-2 mins as previously described (Buchon *et al*, 2009; Clark *et al*, 2015) and washed twice with sterile PBS. The intestine was dissected in sterile PBS with sterile equipment. The intact tissue was carefully taken to prevent loss of lumen contents and malpighian tubules were carefully removed. The dissected intestines were stored at −80°C before analysis. Genomic DNA extraction was performed as described in the manufacturer’s protocol (PowerSoil DNA isolation Kit, MoBio). 15 flies were dissected for each sample to extract DNA. The bacterial load of *Alphaproteobacteria*, *Gammaproteobacteria*, *Bacilli* and total bacteria, different families of bacteria (*Acetobacteraceae*) and *Lactobacili Plantarum* was measured by qPCR with primers listed in Table S1. qPCR results were normalized with *Drosophila* Actin5C as previously described (Claesson *et al*, 2010; Clark *et al*, 2015; Sebald *et al*, 2016).

For colony formation units (CFUs) assays, animals were surface sterilized as described above and 5 intestines were dissected and homogenized in 500 μl PBS. 3 serial dilutions of 1/10 on the homogenates were plated on selective plates for *Acetobacteraceae*, *Lactobacili* and nutrient rich medium (see information for the protocols). The plates were incubated at 29°C and colonies were counted after 48 and 72 hrs as previously described (Broderick *et al*, 2014). At least 2 independent experiments were performed and each with at least 4 biological replicates (n≥4).

### 16S rRNA sequencing

To obtain bacterial genomic DNA (gDNA) from the intestine, flies were surface sterilized before dissection as described above. Whole midguts were maintained as intact tissues to prevent bacterial leakage and the crops of fly guts were removed. 30 guts were dissected for each sample and the gDNA was extracted using PowerSoil DNA isolation kit (MoBio). The gDNA was used to generate amplicons that cover V3 and V4 hypervariable regions of bacteria and Archaea 16S rDNA. Indexed adapters were added to the end of the 16S rDNA amplicons by limited cycle PCR to prepared sequencing library. DNA libraries were multiplexed and sequenced on an Illumina MiSeq instrument.

### Histology

Antibody staining was performed as previously described (Zhou *et al*, 2017). TUNEL staining was performed based on the instruction of the ApopTag Red in situ detection kit (Millipore S7165). Antibodies were used at the following final concentrations: anti-Discs large (1:1200), anti-FasIII (1:1000), anti-beta galactosidase (1:1000), anti-phospho JNK (1:600), anti-pH3 (1:600), anti-Cas3 (1:800), anti-DCP (1:600), All secondary antibodies were diluted at a final concentration of 1:3000. The detail information is provided in Table S3

### Quantification and statistical analysis

For all quantifications, n equals the number of midguts used in the experiment. The results were presented as mean ± SEM. Comparisons between groups were made using the unpaired two-tailed Student’s t test. Statistical analyses for survival curves were performed using the log-rank (Mantel-Cox) test. The significance between groups was expressed as P values. (*) denotes p < 0.05, (**) denotes p < 0.01, (***) denotes p < 0.001, and (ns) denotes no significant difference.

### Quantification of pH3 cell counts per midgut

Mitotic cells marked by phospho-histone 3 staining were counted in the intestine of indicated genotypes. Approximately 10 midguts were dissected and counted in each experiment and data from 3 independent experiments is shown. The results are presented as mean value of cell counts with standard deviation (SD) in the bar figures. Experiments were conducted as described previously (Zhou *et al*, 2017).

## Acknowledgements

We are grateful to M. Funk, F. Heigwer, F. Port and J. Bageritz for comments on the manuscript. We would like to thank B. Edgar, M. Uhlirova, X. Yang, Bloomington *Drosophila* Stock Center and Vienna *Drosophila* Resource Center (VDRC) for fly stocks and antibodies. We thank F. Port for CRISPR lines. We thank K. Han (IPMB, Heidelberg University) and Boutros lab members for helpful discussions and experimental support, especially S. H. Wang, J. Mattila, J. Bageritz, B. Rauscher and A.-L. Boettcher. We would also like to thank the Imaging Core and Genomics and Proteomics Core Facilities at the DKFZ for support. Research in the laboratory of M.B. is supported in part by the DFG Collaborative Research Center SFB873 and the Excellence Cluster CellNetworks.

## Author contributions

J.Z. and M.B. conceived the project. J.Z performed and analyzed experiments. J.Z. and M.B. interpreted experimental data. J.Z. and M.B. wrote the manuscript preparation. M.B. supervised the project.

