## Supplement for "JNK-dependent intestinal barrier failure disrupts host-microbe homeostasis during tumorigenesis"

**SUPPLEMENTAL INFORMATION**

This file includes:

**Supplemental Figures 1-5 (Page 2-10)**

**Supplemental Table 1-3 (Page 11-12)**

**Supplemental Methods (Page 13)**

Recipe of selective plates for bacterial culture

**Supplemental References (Page 13)**

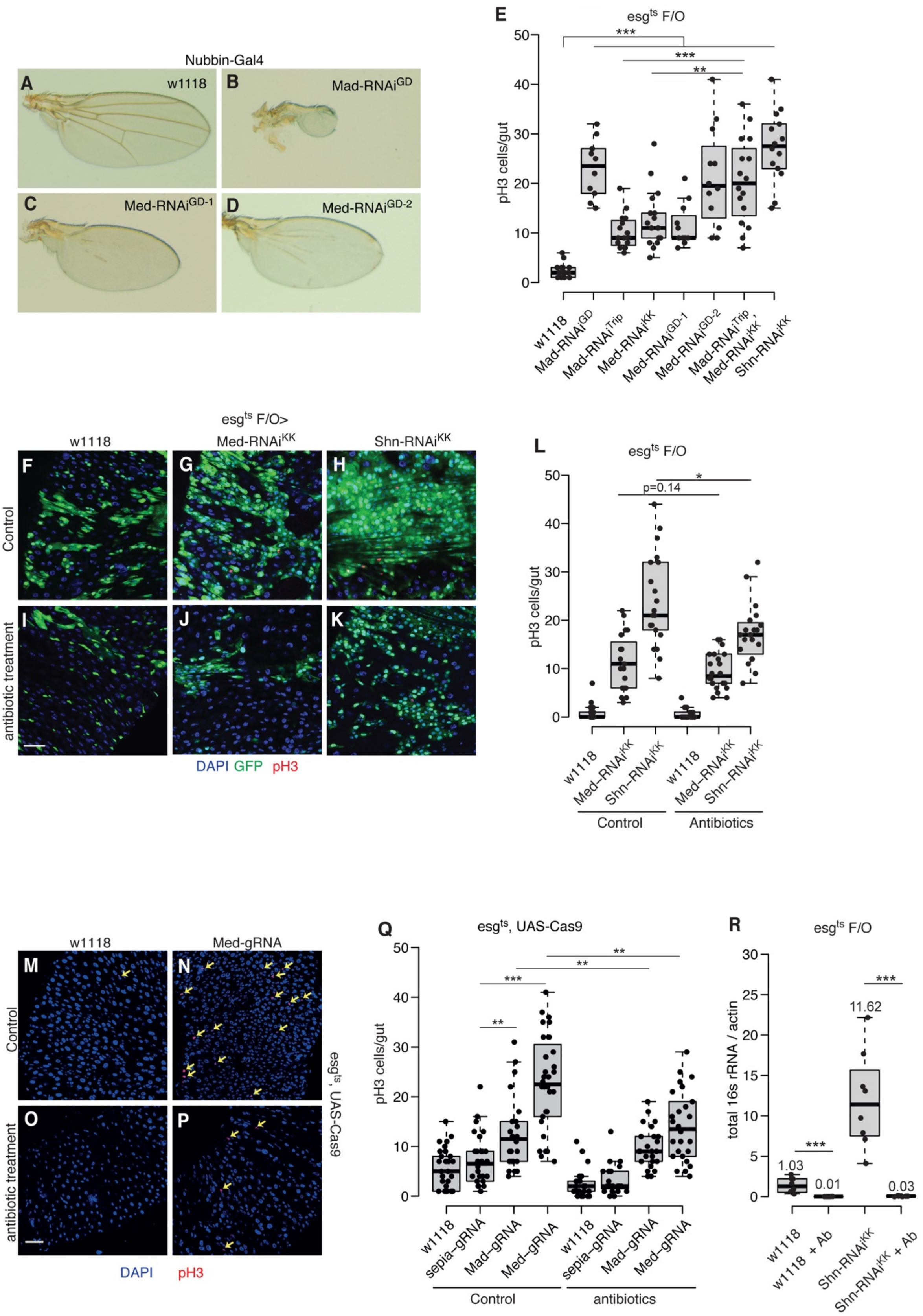

Nubbin-Gal4
A
w1118
B
Mad-RNAi GD
C
Med-RNAi GD-1
D
Med-RNAi GD-2
E
esg ts F/O
pH3 cells/gut
***
***
**
w1118
Mad-RNAi GD
Mad-RNAi Trip
Mad-RNAi KK
Med-RNAi GD-1
Med-RNAi GD-2
Mad-RNAi Trip,
Med-RNAi KK
Shn-RNAi KK
esg ts F/O>
w1118
Med-RNAi KK
Shn-RNAi KK
F
G
H
Control
I
J
K
antibiotic treatment
DAPI GFP pH3
L
esg ts F/O
pH3 cells/gut
p=0.14
*
w1118
Med-RNAi KK
Shn-RNAi KK
Control
Antibiotics
w1118
Med-gRNA
M
N
Control
O
P
antibiotic treatment
esg ts, UAS-Cas9
DAPI
pH3
Q
esg ts, UAS-Cas9
pH3 cells/gut
**
***
**
**
w1118
sepia-gRNA
Mad-gRNA
Med-gRNA
Control
antibiotics
R
esg ts F/O
total 16s rRNA / actin
1.03
0.01
11.62
0.03
***
***
w1118
w1118 + Ab
Shn-RNAi KK
Shn-RNAi KK + Ab

**Figure S1. Intestinal microbiota triggers tumor growth.**
(A-D) Wing-specific Nubbin-Gal4 (*Nub-Gal4*) driven Mad-RNAi$^{GD}$ (B), Med-RNAi$^{GD-1}$ (C) and Med-RNAi$^{GD-2}$ (D) show vein formation and growth defect during development similar to Dpp loss of function phenotype, compared to control (A).
(E) Quantification of pH3-positive cells per adult midgut of the indicated genotypes after 7 days at 29°C.
(F-K) Representative images of the posterior midgut of *esg$^{ts}$F/O>w1118* (F, I), *esg$^{ts}$F/O>Med-RNAi$^{KK}$* (G, J) and *esg$^{ts}$F/O>Shn-RNAi$^{KK}$* (H, K) flies in standard and antibiotics-treated fly food, stained with pH3 antibody in red. *Act-GFP* is shown in GFP, nuclei are stained with DAPI (blue).
(L) Quantification of pH3-positive cells per adult midgut of the indicated genotypes in standard and antibiotics-treated fly food after 7 days at 29°C. Med-RNAi only led to slight but non-significant decrease in stem cell proliferation upon antibiotic feeding (Fig EV1L, O, Q), which could be caused by the relatively mild phenotype with Med-RNAi (Zhou et al., 2015).
(M-P) Representative images of the posterior midgut of *esg$^{ts}$, UAS-Cas9>w1118*
(M, O), *esg$^{ts}$, UAS-Cas9>Med-gRNA* (N, P) flies in standard and antibiotics-treated fly food at 18°C for 30 days, stained with pH3 antibody in red, nuclei are stained with DAPI (blue), *esg>GFP* is not observed since only temporal sgRNAs/Cas9 expression is driven by *esg$^{ts}$*.
(Q) Quantification of pH3-positive cells per adult midgut of the indicated genotypes in standard and antibiotics-treated fly food for 30 days at 18°C after inducing CRISPR/Cas9 mediated mutagenesis.
(R) Bacterial level in the intestine of *esg$^{ts}$F/O>Shn RNAi* and *esg$^{ts}$F/O>w1118* with or without antibiotic treatment, assayed by qPCR of 16s rRNA gene at 29°C for 8 days.* $p<0.05$; ** $p<0.01$; *** $p<0.001$). Scale bars: 30 µm (F-K, M-P).

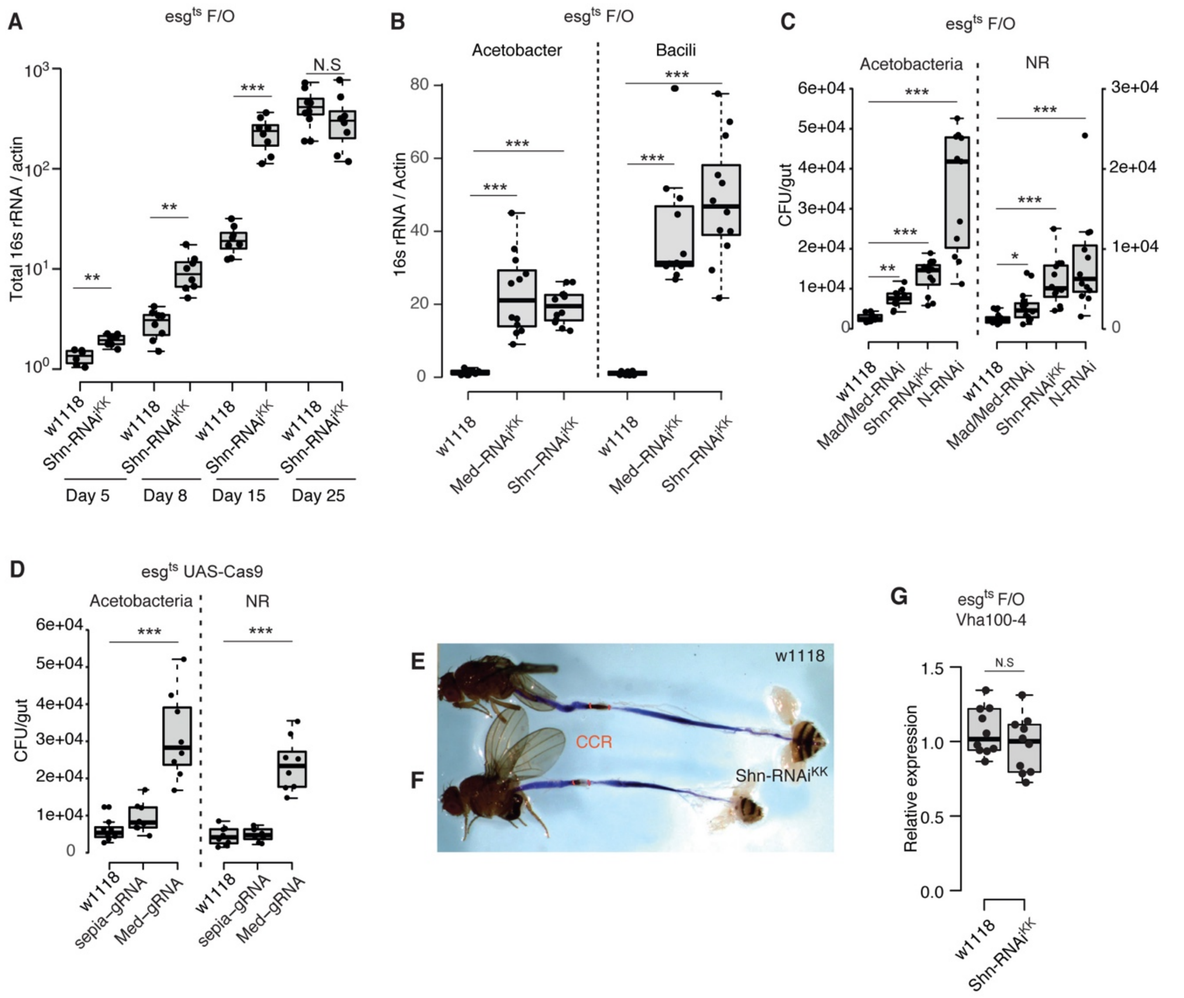


**Figure S2.** Intestinal tumorigenesis induces microbiota dysbiosis.

(A) Bacterial levels in the intestine of *$esg^{ts}$F/O>Shn-RNAi$^{KK}$* and *$esg^{ts}$F/O>w1118* flies, assayed by qPCR of 16s rRNA gene at 5, 8, 15 and 25 days at 29°C. See Table EV1 for further information on bacterial qPCR.

(B) *Alphaproteo*-bacteria and *Bacili*-bacteria levels in the intestine of *$esg^{ts}$F/O>Shn RNAi*,*$esg^{ts}$F/O>Med RNAi* and *$esg^{ts}$F/O>WT* at 15 days, assayed by taxa-specific qPCR of the 16s rRNA gene. n=3 replicates of 15 dissected intestines.

(C) Assay of colony-forming units (CFUs) in the dissected intestine of *$esg^{ts}$F/O>Mad-RNAi$^{Trip}$/Med-RNAi$^{KK}$, $esg^{ts}$F/O>Shn-RNAi$^{KK}$, $esg^{ts}$F/O>Notch (N)-RNAi* and *$esg^{ts}$F/O>w1118* genotypes after 15 days at 29°C.

(D) Assay of colony-forming units (CFUs) in the dissected intestine of *$esg^{ts}$;UAS-Cas9>Med-gRNA, $esg^{ts}$;UAS-Cas9>sepia-gRNA* and *$esg^{ts}$;UAS-Cas9>w1118* genotypes at 18°C for 30 days after inducing mutagenesis. Homogenates of intestines from flies at 15 days were plated on selective plates for *Acetobacteria* or on nutrient-rich medium plate.

(E-F) Gastrointestinal tract of *$esg^{ts}$ F/O>w1118* (E) and *$esg^{ts}$F/O> Shn-RNAi$^{KK}$* (F) flies fed with the PH indicator Bromophenol blue.

(G) RT-qPCR analysis of the intestine shows no difference in the expression level of *Vha100-4* gene in Shn depleted midgut.* p<0.05; ** p<0.01; *** p<0.001.

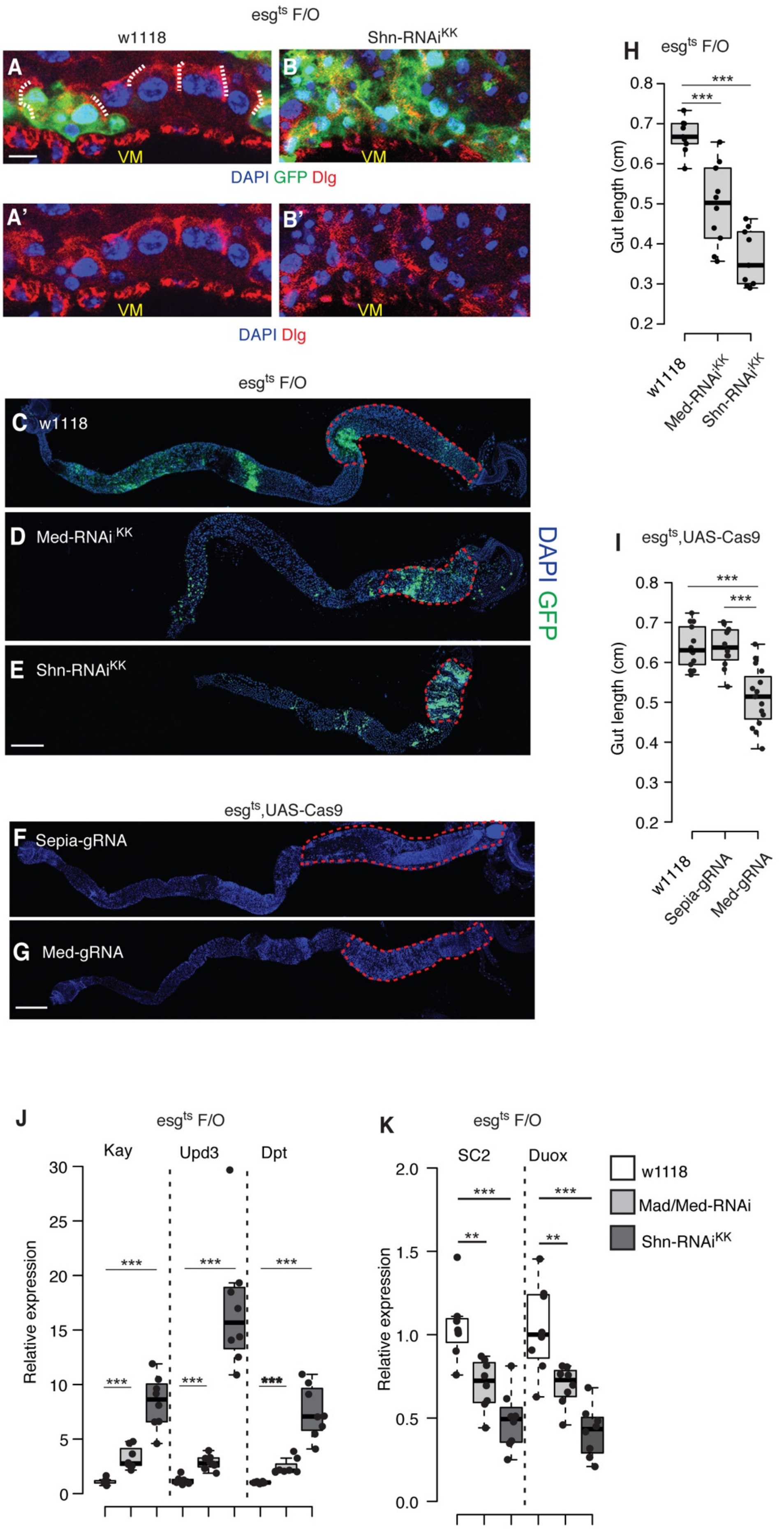
esgts F/O
w1118
Shn-RNAiKK
A
B
VM
DAPI GFP Dlg
A'
B'
DAPI Dlg
esgts F/O
C w1118
D Med-RNAi KK
E Shn-RNAiKK
DAPI GFP
esgts,UAS-Cas9
F Sepia-gRNA
G Med-gRNA
H esgts F/O
Gut length (cm)
w1118
Med-RNAiKK
Shn-RNAiKK
I esgts,UAS-Cas9
Gut length (cm)
w1118
Sepia-gRNA
Med-gRNA
J esgts F/O
Kay
Upd3
Dpt
Relative expression
K esgts F/O
SC2
Duox
Relative expression
w1118
Mad/Med-RNAi
Shn-RNAiKK

**Figure S3.** Tumor growth impairs intestinal barrier function.

(A-B) Higher magnification transversal view of the posterior midgut epithelium of *esg*$^{ts}$*F/O>w1118* (L), *esg*$^{ts}$*F/O>Shn-RNAi*$^{KK}$ (M) flies at 29°C for 7 days, stained with anti-Disc large antibody.

(C-E) Representative images of the midgut of *esg*$^{ts}$*F/O>w1118* (C), *esg*$^{ts}$*F/O>Med-RNAi*$^{KK}$ (D), *esgF/O>Shn-RNAi*$^{KK}$ (E) flies at 29°C for 15 days. *Act-GFP*is shown in GFP, nuclei are stained with DAPI (blue). The posterior midgut region is indicated by a red dashed line. Note that the preference of tumor initiation and growth in certain regions of the intestine of Med or Shn RNAi flies (regions R2, R4 and R5; Fig EV3D and E) is similar as described in loss-of Notch induced tumors (Marianes and Spradling, 2013).

(F-G) Representative images of the midgut of *esg*$^{ts}$*; UAS-Cas9 > Med-gRNA, esg*$^{ts}$*; UAS-Cas9 > sepia-gRNA*. The posterior midgut region is indicated by a red dashed line.

(H) Quantification of the length of intestine of *esg*$^{ts}$*F/O>Shn-RNAi*$^{KK}$, *esg*$^{ts}$*F/O>Med-RNAi*$^{KK}$ and *esg*$^{ts}$*F/O>w1118* at 29°C for 15 days.

(I) Quantification of the length of intestine of *esg*$^{ts}$*;UAS-Cas9>Med-gRNA, esg*$^{ts}$*;UAS-Cas9>sepia-gRNA* and *esg*$^{ts}$*;UAS-Cas9>w1118* genotypes at 18°C for 30 days after inducing mutagenesis.

(J) RT-qPCR analysis of the intestine shows that the expression levels of JNK (*Kay*), Relish (*Dpt*) and regenerative cytokine (Upd3) pathway component genes are increased in *esg*$^{ts}$*F/O>Mad-RNAi*$^{Trip}$*/Med-RNAi*$^{KK}$ and *esg*$^{ts}$*F/O>Shn-RNAi*$^{KK}$ midgut at 29°C for 7 days.

(K) RT-qPCR analysis of the intestine shows that the expression levels of Relish (*PGRP-SC2*:*SC2*) and ROS production (*Duox*) pathway component genes are decreased in *esg*$^{ts}$*F/O>Mad-RNAi*$^{Trip}$*/Med-RNAi*$^{KK}$ and *esg*$^{ts}$*F/O>Shn-RNAi*$^{KK}$ midgut at 29°C for 7 days. * p<0.05; ** p<0.01; *** p<0.001. Scale bars: 10 µm (A-B), 250 µm (C-G).

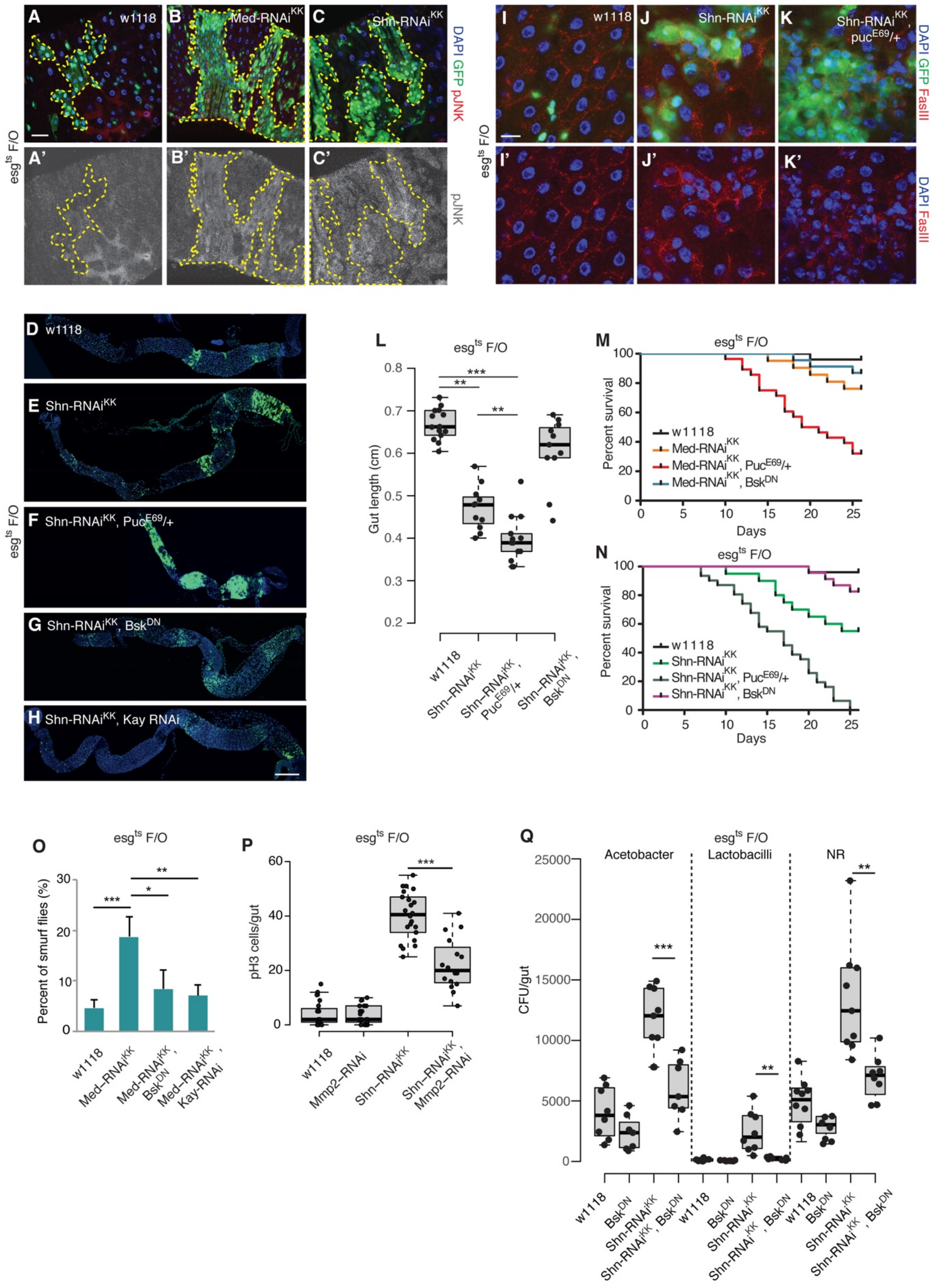

A w1118
B Med-RNAi^KK
C Shn-RNAi^KK
DAPI GFP pJNK
A' B' C' pJNK
esg^ts F/O
I w1118
J Shn-RNAi^KK
K Shn-RNAi^KK, puc^E69/+
DAPI GFP FasIII
I' J' K' DAPI FasIII
D w1118
E Shn-RNAi^KK
F Shn-RNAi^KK, Puc^E69/+
G Shn-RNAi^KK, Bsk^DN
H Shn-RNAi^KK, Kay RNAi
L esg^ts F/O
Gut length (cm)
w1118
Shn-RNAi^KK
Shn-RNAi^KK, Puc^E69/+
Shn-RNAi^KK, Bsk^DN
M esg^ts F/O
Percent survival
Days
w1118
Med-RNAi^KK
Med-RNAi^KK, Puc^E69/+
Med-RNAi^KK, Bsk^DN
N esg^ts F/O
Percent survival
Days
w1118
Shn-RNAi^KK
Shn-RNAi^KK, Puc^E69/+
Shn-RNAi^KK, Bsk^DN
O esg^ts F/O
Percent of smurf flies (%)
w1118
Med-RNAi^KK
Med-RNAi^KK, Bsk^DN
Med-RNAi^KK, Kay-RNAi
P esg^ts F/O
pH3 cells/gut
w1118
Mmp2-RNAi
Shn-RNAi^KK
Shn-RNAi^KK, Mmp2-RNAi
Q esg^ts F/O
CFU/gut
Acetobacter
Lactobacilli
NR
w1118
Bsk^DN
Shn-RNAi^KK
Shn-RNAi^KK, Bsk^DN

**Figure S4.** Tumor leads to intestinal JNK signaling activation

(A-C) Immunofluorescence images of midguts of *esg*$^{ts}$*F/O>w1118* (A), *esg*$^{ts}$*F/O>Med-RNAi* (B) and *esg*$^{ts}$*F/O>Shn-RNAi* (C) with phospho-JNK (pJNK) staining at 29°C for 7 days.

(D-H) Representative image of intestines from *esgF/O>w1118* (D), *esg*$^{ts}$*F/O>Shn-RNAi* (E), *esg*$^{ts}$*F/O>Shn-RNAi*;*Puc*$^{E69}$/+ (F) and *esg*$^{ts}$*F/O>Shn-RNAi*;*Bsk*$^{DN}$ (G), *esg*$^{ts}$*F/O>Shn-RNAi*; *Kay-RNAi* (H) after 7 days RNAi induction at 29°C.

(I-K) Representative images of the posterior midgut of *esg*$^{ts}$*F/O>w1118* (I), *esg*$^{ts}$*F/O>Shn-RNAi* (J), *esg*$^{ts}$*F/O>Shn-RNAi*, *Puc*$^{E69}$/+ (K) flies at 29°C for 7 days, stained with anti-FasIII antibody.

(L) Quantification of the length of intestine of indicated genotypes at 29°C for 15 days.

(M-N) Med RNAi (M) or Shn RNAi (N) induced intestinal tumor decreases survival of flies compared to WT. Introducing heterozygous mutation of Puckered (*Puc*$^{E69}$) enhances the phenotype. Conversely, suppressing JNK by expressing Bsk$^{DN}$ or Kay RNAi rescues Med or Shn loss induced organismal death.

(O) Smurf assay for gut permeability of indicated genotype at 29°C for 25 days. High percentage of Smurf flies are shown in the group of Med RNAi flies and JNK suppression (Bsk$^{DN}$ or Kay RNAi) reduces Med RNAi related Smurf phenotype.

(P) Quantification of pH3-positive cells per adult midgut of the indicated genotypes at 29°C for 10 days.

(Q) The assay of colony-forming units (CFUs) in dissected intestine of *esg*$^{ts}$*F/O>UAS-Bsk*$^{DN}$*, esg*$^{ts}$*F/O>Shn-RNAi*$^{KK}$*, esg*$^{ts}$*F/O>Shn-RNAi*$^{KK}$*&UAS-Bsk*$^{DN}$ and *esg*$^{ts}$*F/O>WT* at 29°C for 15 days. The homogenates of intestine from flies at 15 days were plated on selective plates for *Acetobacteria*, *Lactobacilli* (MRS Agar) or on Nutrient-rich medium plate. Scale bars: 30 µm (A-C), 250 µm (D-H),10 µm (I-K)

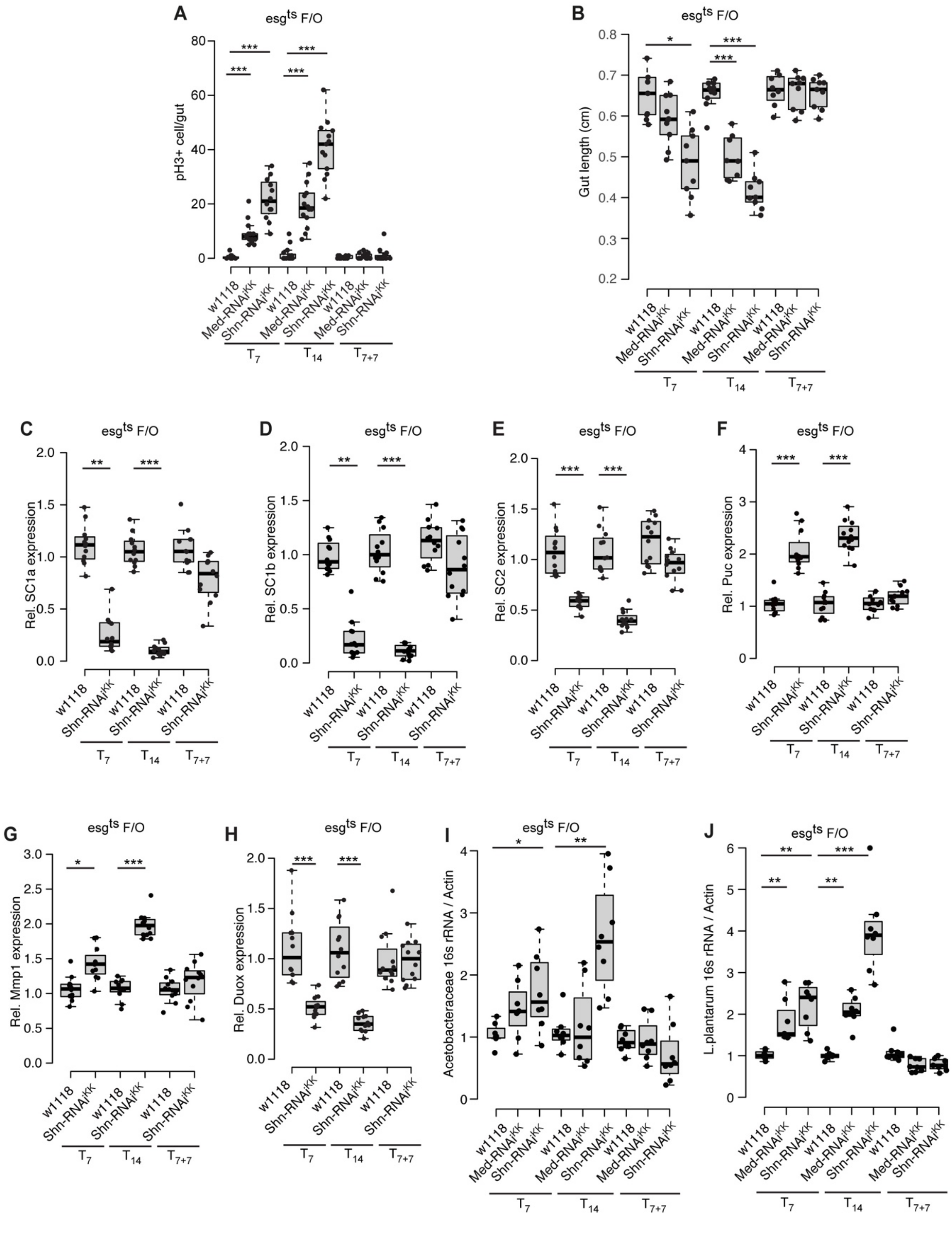

A
esg$^{ts}$ F/O
pH3+ cell/gut
B
Gut length (cm)
C
Rel. SC1a expression
D
Rel. SC1b expression
E
Rel. SC2 expression
F
Rel. Puc expression
G
Rel. Mmp1 expression
H
Rel. Duox expression
I
Acetobacteraceae 16s rRNA / Actin
J
L.plantarum 16s rRNA / Actin
w1118
Med-RNAi$^{KK}$
Shn-RNAi$^{KK}$
$T_7$
$T_{14}$
$T_{7+7}$

**Figure S5.** The intestinal barrier dysfunction is associated with tumor related dysbiosis.
(A) Quantification of pH3-positive cells per adult midgut of the indicated genotypes.
(B) Quantification of intestinal length of of *esg*$^{ts}$*F/O>Shn-RNAi*, *esg*$^{ts}$*F/O>Med-RNAi* and *esg*$^{ts}$*F/O>w1118* in tumor ‘start’, ‘progression’ and ‘recovery’ conditions using NeuronJ, an ImageJ plugin. The number shown in the box indicates the number of intestines used in the experiment. The significant differences in the length of intestine between RNAi group (*esg*$^{ts}$*F/O >Shn-RNAi* or *Med-RNAi*) and the control group (*esg*$^{ts}$*F/O>w1118*) are indicated by *p*value.
(C-H) RT-qPCR analysis of the intestine shows that the expression levels of Relish (*PGRP-SC1a*-C, *PGRP-SC1b*-D, *PGRP-SC2*:*SC2*-E,), JNK (*Puc*-F, *Mmp1*-G) and ROS production (*Duox*-H) pathway component genes are varied in *esg*$^{ts}$*F/O>Shn-RNAi* midgut in the conditions of tumor start (*Shn-RNAi*$^{KK}$ compared to *WT* at 29°C for 7 days), tumor progression (*Shn-RNAi*$^{KK}$ compared to *WT* at 29°C for 14 days) and tumor recovery (*Shn-RNAi*$^{KK}$ compared to *WT* at 29°C for 7 days, then shift to 18°C for 7 days).
(I-J) Bacterial levels in the intestine of *esg*$^{ts}$*F/O>Shn RNAi*, *esg*$^{ts}$*F/O>Med RNAi* and *esg*$^{ts}$*F/O>w1118* during tumor start, progression and recovery, assayed by taxon-specific qPCR of 16s rRNA gene. (I) *Acetobacteriaceae*, (J) *Lactobacilli*. * p<0.05; ** p<0.01; *** p<0.001.

**Supplemental Table 1:** Sequence information for Bacterial qPCRs

| Name | Primer direction | Sequences | References |
|---|---|---|---|
| ***Alpha*** | Left | CCAGGGCTTGAATGTAGAGGC | 1 |
| ***Alpha*** | Right | CCTTGCGGTTCGCTCACCGGC | 1 |
| ***Bacilli*** | Left | CGACCTGAGAGGGTAATCGGC | 1 |
| ***Bacilli*** | Right | CGACCTGAGAGGGTAATCGGC | 1 |
| **Gamma** | Left | GGTAGCTAATACCGCATAACG | 1 |
| **Gamma** | Right | TCTCAGTTCCAGTGTGGCTGG | 1 |
| ***Universal*** | Left | AGAGTTTGATCCTGGCTCAG | 1 |
| ***Universal*** | Right | CTGCTGCCTYCCGTA | 1 |
| ***Acetobacter*** | Left | CZAGTGTAGAGGTGAAATT | 2 |
| ***Acetobacter*** | Right | CCCCGTCAATTCCTTTGAGTT | 2 |
| ***L.plantarum*** | Left | TGATCCTGGCTCAGGACGAA | 2 |
| ***L.plantarum*** | Right | TGCAAGCACCAATCAATACCA | 2 |
| ***Drosophila* Actin5C** | Left | TTGTCTGGGCAAGAGGATCAG | 1 |
| ***Drosophila* Actin5C** | Right | ACCACTCGCACTTGCACTTTC | 1 |

**Supplemental Table 2:** Sequence information for qPCRs

| Name | Primer direction | Sequences | Probe No. |
|---|---|---|---|
| AttD | Left | gtttatggagcggtcaacg | 44 |
| AttD | Right | tctggaagagattggcttgg | 44 |
| Dpt | Left | cacgagattggactgaatgg | 56 |
| Dpt | Right | tttccagctcggttctgagt | 56 |
| Duox | Left | acgtgtccacccaatcgcacgag | SyBR green |
| Duox | Right | aagggtggtggtccagtcagtcg | SyBR green |
| InR | Left | tacaagtgcgggcgattc | 141 |
| InR | Right | ttcacgtgatctcaatcatgc | 141 |
| Kay | Left | cagcatcagcgacaggatta | 113 |
| Kay | Right | tctggccggtctcaaagtt | 113 |
| Mmp1 | Left | ccgatttgctgttgactcg | 60 |
| Mmp1 | Right | tgtatccgcgtctcttaaagc | 60 |
| Mmp2 | Left | ggcaaaaaccgctatctcc | 113 |
| Mmp2 | Right | cagagcacccgtttccag | 113 |
| Pi3k21B | Left | cagatctatatccattgattctttcg | 89 |
| Pi3k21B | Right | gccgcatcgtcgagtagt | 89 |
| PGRP-SC1a | Left | ggcaactacctcagctacgc | 138 |
| PGRP-SC1a | Right | acgggtctcgcagtaggag | 138 |
| PGRP-SC1b | Left | aaagtggttacagcccaacg | 56 |

| | | | |
|---|---|---|---|
| PGRP-SC1b | Right | aagtgcagctcgaaagacct | 56 |
| PGRP-SC2 | Left | ccaagtctatcggcatctcc | 61 |
| PGRP-SC2 | Right | gagcagaggtgagggtgttg | 61 |
| Puc | Left | cgtcatcatcaacggcaat | 70 |
| Puc | Right | aggcggggtgtgtttctat | 70 |
| Rp49 | Left | ttccttgacgtgccaaaact | 159 |
| Rp49 | Right | aatgatctataacaaaatcccctga | 159 |
| Socs36E | Left | aaaaagccagcaaaccaaaa | 43 |
| Socs36E | Right | aggtgatgacccattggaag | 43 |
| spitz | Left | gcgggtgtttttgttgtcat | 31 |
| spitz | Right | ttggaatcgggtttctctaca | 31 |
| Upd2 | Left | aagttcctgccgaacatgac | 44 |
| Upd2 | Right | atccttgcggaacttgtactg | 44 |
| Upd3 | Left | cccagccaacgatttttatg | 165 |
| Upd3 | Right | tgttaccgctccggctac | 165 |

**Supplemental Table 3:** The detail information for antibodies used in this study

| **Antibodies** | **Source** | **Order information** |
|---|---|---|
| mouse monoclonal anti-discs large | DSHB | Cat# 4F3; RRID: AB_528203 |
| mouse monoclonal anti-FasIII | DSHB | Cat# 7G10; RRID: AB_528238 |
| rabbit polyclonal anti-beta galactosidase | DSHB | Cat# 40-1a; RRID: AB_2314509 |
| rabbit polyclonal anti-phospho JNK | Millipore | Cat# 07-175; RRID: AB_310412 |
| rabbit polyclonal anti-phospho-Histone 3 (Ser10) | Millipore | Cat# 06-570; RRID: AB_310177 |
| Rabbit anti-cleaved caspase 3 | Millipore | Cat# AB3623, RRID:AB_91556 |
| Rabbit anti-cleaved Drosophila Dcp-1 (Asp216) | Millipore | Cat# AB9578, RRID:AB_2721060 |
| Goat anti-rabbit IgG (H+L) secondary antibody Alexa 594 | Fisher Scientific | Cat# 35561, RRID:AB_1965951 |
| Goat anti-mouse IgG (H+L) secondary antibody Alexa 594 | Fisher Scientific | Cat# 35511, RRID:AB_1965950 |
| Goat anti-rabbit IgG (H+L) secondary antibody Alexa 633 | Fisher Scientific | Cat# 35513, RRID:AB_1965952 |
| Goat anti-mouse IgG (H+L) secondary antibody Alexa 633 | Fisher Scientific | Cat# 35563, RRID:AB_1965953 |

**Supplemental Methods**

*Recipe of selective plates for bacterial culture*
*Acetobacteriaceae*: 25 g/l D-mannitol, 5 g/l yeast extract, 3 g/l peptone, and 15 g/l agar.
*Lactobacilli* MRS agar: 70 g/l BD Difco Lactobacilli MRS agar.
Nutrient Rich Broth: 23 g/l BD Difco Nutrient agar.
All medium were autoclaved at 121°C for 15 min.
